# TRIB1 modulates transcriptional programming in breast cancer cells to regulate cell proliferation

**DOI:** 10.1101/2023.07.06.547928

**Authors:** Hamish D. McMillan, Evangelia K. Papachristou, Jody Hazlett, Soleilmane Omarjee, Jason S. Carroll, Michael A. Black, Peter D. Mace, Anita K. Dunbier

## Abstract

The pseudokinase Tribbles Homolog 1 (*TRIB1*) is a known driver of tumorigenesis in acute myeloid leukemia and is encoded upstream of the oncogene *MYC* at the 8q24 locus. We observed that *TRIB1/MYC* co-amplification is associated with decreased relapse-free and overall survival in breast cancer patients, but the role of *TRIB1* in this disease has not been well characterized. *TRIB1* knockdown in multiple breast cancer cell lines inhibited cell proliferation and suppressed *MYC* expression, implicating *TRIB1* in breast cancer cell proliferation. Transcriptomic and cell cycle analysis revealed cell cycle regulation as the likely mechanism through which *TRIB1* influences breast cancer cell proliferation. *TRIB1* knockdown also resulted in significant changes in both estrogen receptor (ER) and β-catenin associated transcription. Interrogating the TRIB1 interactome in breast cancer cells by qPLEX-RIME reinforced the known association between TRIB1 and ubiquitination, while revealing a range of previously undescribed TRIB1 associated factors. Further analysis of the association between TRIB1, β-catenin and FERMT2 suggests TRIB1 may regulate β-catenin activity by controlling the levels of both β-catenin, and its co-factor FERMT2. Together, these results suggest that coregulation of β-catenin and ER-driven transcription by TRIB1, facilitates regulation of *MYC* expression and breast cancer cell proliferation.

**Significance:** The pseudokinase *TRIB1* is frequently co-amplified in breast cancers with the potent oncogene *MYC*, although the functional consequences of this event are not well understood. This study demonstrates *TRIB1* is a regulator of cell cycle progression and *MYC* expression in breast cancer cells. It also profiles *TRIB1*-associated proteins in breast cancer cells, demonstrating conservation of TRIB1’s canonical interaction with COP1 and reveals associations with members of the wider ubiquitination machinery, a range of transcriptional regulators and chromatin remodelers. The data presented provide insight into the function of TRIB1 in breast cancer and the role of TRIB1 in transcriptional regulation.

## Introduction

Tribbles homolog 1 (*TRIB1*) encodes the leukaemia oncoprotein, pseudokinase TRIB1 (1, 2). During myeloid differentiation, *TRIB1* overexpression leads to the degradation of the transcription factor C/EBPα, preventing myeloid differentiation and driving development of acute myeloid leukaemia (3–6). Degradation of C/EBPα occurs via COP1 ubiquitin ligase, and recruitment of substrates to COP1 by TRIB1 has been extensively characterized at both the structural and cellular level (3–10). Despite extensive characterization of *TRIB1* in leukaemia, the importance of *TRIB1* in solid tumors is less well-understood. Nonetheless, various studies have found *TRIB1* to be associated with the development and progression of several solid tumor types including breast cancer (11, 12), hepatocellular carcinoma (13), glioma (14), gastric cancer (15) and prostate cancer (16, 17).

*TRIB1* is co-located within a frequently amplified chromosome region 8q24, approximately 2 MB upstream of the potent oncogene *MYC* (18). *MYC* is overexpressed or deregulated in approximately 70% of all cancers (19), and amplification of the 8q24 amplicon is a common mechanism of *MYC* dysregulation. A functional association between *TRIB1* expression and *MYC* overexpression has been established, with *TRIB1* knockdown being identified as synthetic lethal to cells with exogenous *MYC* overexpression (20), however the nature of this relationship in the endogenous amplification context is poorly described. The genomic, and potential functional relationship, between *TRIB1* and *MYC* suggest further investigation of the role of *TRIB1* in solid tumors is warranted.

A number of studies have investigated the role of *TRIB1* in immune (21–25), hepatic (26–28) and adipose tissues (29, 30). However, relatively little is known about the functional role of TRIB1 in breast cancer, despite the frequent co-amplification with *MYC*. Assessment of the *TRIB1* knockdown phenotype, combined with parallel transcriptomic and proteomic analyses, was used to investigate the role of TRIB1 in breast cancer cells. Integrated examination of the TRIB1 interactome with the transcriptional and phenotypic consequences of *TRIB1* knockdown, in breast cancer cells, reveal that TRIB1 exists in complex with key breast cancer transcription factors, regulating cell proliferation in concert with *MYC*.

## Results

### *TRIB1/MYC* co-amplification is associated with decreased survival in breast cancer

*TRIB1* and *MYC* are significantly co-amplified (cBioPortal log_2_OR > 3, *P* < 0.001 (31)) in breast cancer patients in both the METABRIC (32, 33) and TCGA (34) datasets, yet the role of *TRIB1* in amplification-driven *MYC* overexpression is not known. Interrogation of the METABRIC breast cancer dataset revealed that approximately 23% of breast tumors carry *TRIB1/MYC* co-amplification. Only 2.6% of tumors carry *MYC* amplification without *TRIB1* amplification, and 0.5% of tumors carry *MYC*-independent *TRIB1* amplification (31–33). *TRIB1/MYC* co-amplification is significantly associated with decreased relapse-free (Fig. 1A) and overall survival (Supplementary Fig. 1A) in a subtype independent manner. The strong association between *TRIB1* and *MYC* copy number observed in patient samples (Supplementary Fig. 1B) is recapitulated in breast cancer cell lines (Fig. 1B).

**Figure 1:**
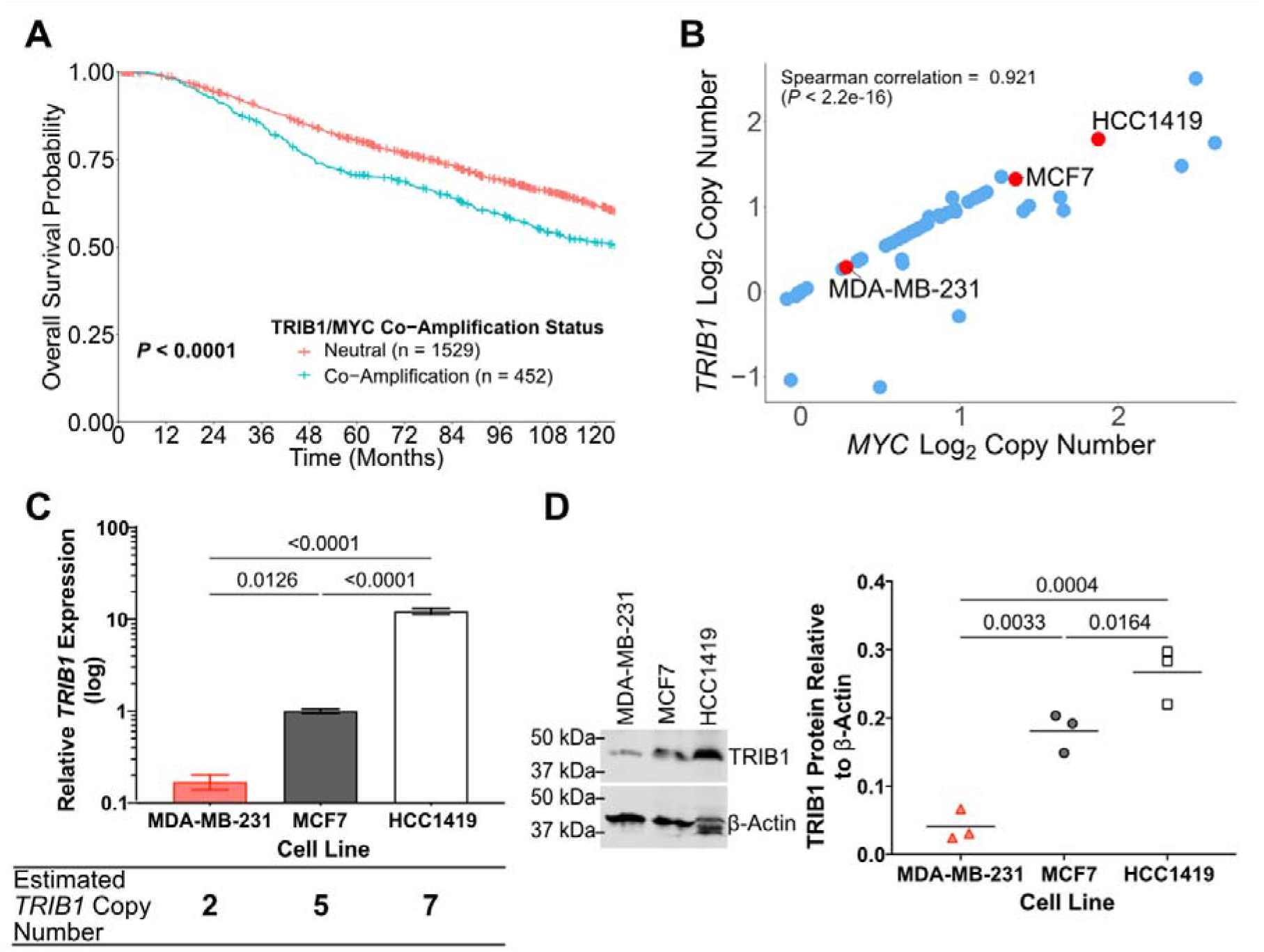
*TRIB1/MYC* co-amplification is associated with decreased long term survival and is recapitulated in breast cancer cell lines. **A**, Kaplan-Maier overall survival plot of METABRIC data, stratified based on *TRIB1/MYC* co-amplification status (*31–33*). **B**, Comparison of estimated *TRIB1* and *MYC* copy number in the breast cancer cell lines from the CCLE determined from Affymetrix SNP 6.0 array data (GSE36138) (*35, 36*). **C**, *TRIB1* expression in MDA-MB-231 (red), MCF7 (grey) and HCC1419 (white) cells was determined by RT-qPCR after 72 hours of growth and normalized to the geometric mean of *PUM1* and *FKBP15* expression. All data are the mean ± SD of three biological replicates (each biological replicate is the average of three technical replicates). Significance calculated using multiple comparison two-way ANOVA with Tukey *P-value* adjustment. Absolute copy number estimated from CCLE data (Fig. 1B) by taking the inverse of the log_2_CN and multiplying by two. **D**, Representative western blot of TRIB1 levels in MDA-MB-231, MCF7 and HCC1419 cells after 72 hours of growth. The gel was loaded with 50 μg of total protein from each cell line. TRIB1 densitometry performed on three biological replicates (second and third replicates supplementary Fig. 1E) and normalized to β-actin. Significance determined using multiple comparison one-way ANOVA analysis with Holm-Šídák *P-value* adjustment.

To investigate the role of *TRIB1* in breast cancer, three breast cancer cell lines with varying *TRIB1* copy number were selected from the Cancer Cell Line Encyclopedia (CCLE) breast cancer panel (Fig. 1B). Cell lines representing three breast cancer subtypes were selected for further investigation: triple negative MDA-MB-231; estrogen receptor positive (ER+) MCF7; and HER2-positive HCC1419. The relative expression of *TRIB1*, as determined by CCLE transcriptional analysis, suggested *TRIB1* copy number directly influences *TRIB1* expression (Fig. 1B, Supplementary Fig. 1C).

The association between copy number and expression level was confirmed in the selected cell lines using qPCR analysis, which showed mRNA expression increased proportionately with increasing estimated copy number (Fig. 1B and Fig. 1C). The MDA-MB-231 cells have the lowest levels of *TRIB1* expression, with the MCF7 and HCC1419 having six-fold and 73-fold greater expression, respectively (Fig. 1C). A similar pattern across the cell lines is seen at the protein level, with the three cell lines having significantly different TRIB1 protein levels (Fig. 1D). TRIB1 protein levels are less variable than transcript levels, with only a 6.5-fold difference between MDA-MB-231 and HCC1419 TRIB1 protein levels (Fig. 1D). Taken together, these results suggest that *TRIB1* amplification leads to higher levels of *TRIB1* transcript and protein in breast cancer cells. This trend was also observed at the transcript level in breast cancer primary tumors from the METABRIC study (Supplementary Fig. 1D).

### Knockdown of *TRIB1* expression inhibits breast cancer cell proliferation and modulates *MYC* gene expression

The role of *TRIB1* in breast cancer cell proliferation was evaluated by assessing the effect of *TRIB1* knockdown on the proliferation of the three selected cell lines. *TRIB1* knockdown was achieved by transfection of cells with siRNA SMARTpool targeting *TRIB1*, and cell proliferation was assessed by IncuCyte live cell imaging and nuclei counting. *TRIB1* knockdown in MCF7 cells (Supplementary Fig. 2A) was associated with a 1.9-fold (*P* = 0.0011) decrease in the rate of change in confluency, as measured by live cell imaging confluency tracking (Fig. 2A, Supplementary Fig. 3A-D). *TRIB1* knockdown by siRNA also led to a 2.3-fold (*P* = 0.005) decrease in cell number 72 hours post transfection (Fig. 2B). This phenotype is recapitulated by shRNA mediated knockdown in MCF7 cells, where *TRIB1* resulted in a 1.8-fold (Holm-Šídák adj*.p-value* = 0.039, multiple unpaired T-tests) decrease in cell number 96 hours after seeding (Fig. 2C).

**Figure 2:**
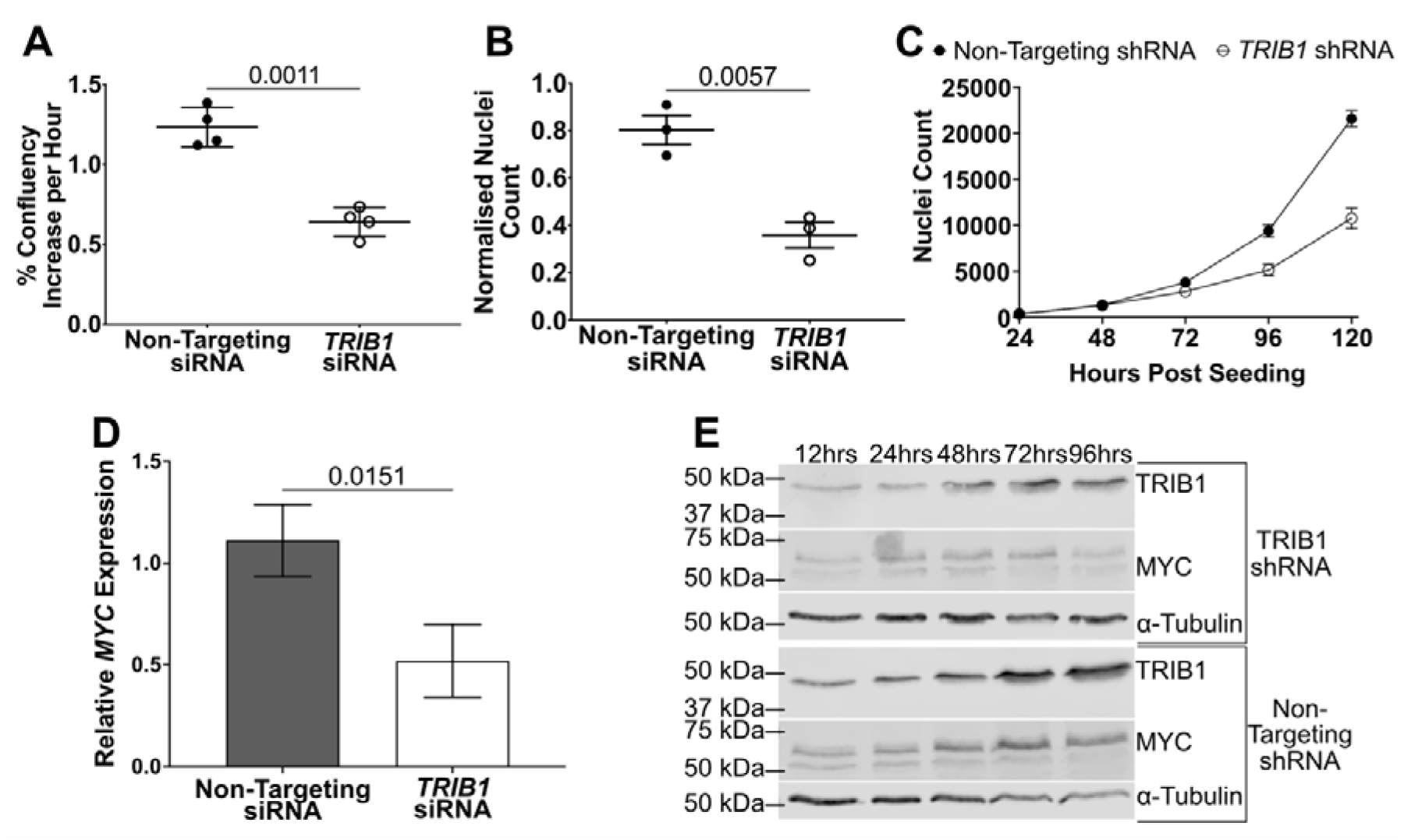
*TRIB1* knockdown slows the proliferation of MCF7 cells and supresses the expression of *MYC*. **A**, Rate of change in confluency of MCF7 cells after transfection with the *TRIB1* or Non-Targeting ON-TARGETplus siRNA SMARTpools. Growth rate determined from IncuCyte live cell imaging confluency traces collected for 168 hours post-transfection (Supplementary Fig. 3A-D). Data are the mean ± SD of four biological replicates with significance determined by paired T-test. **B**, MCF7 nuclei count 72 hours after transfection with *TRIB1* or Non-Targeting ON-TARGETplus siRNA SMARTpools, normalized to untreated nuclei count. Data are the mean ± SEM of three biological replicates, each containing three technical replicates. Significance determined by unpaired T-test. **C**, Nuclei counts of MCF7 monoclonal cell lines with *TRIB1* or non-targeting shRNA expression induced with 2 μg/mL doxycycline. Each point is the mean ± SEM from three biological replicates, each the mean of six technical replicates. **D**, *MYC* expression, in MCF7 cells, 72 hours after siRNA SMARTpool transfection. Expression was determined by RT-qPCR and made relative to the geometric mean of *FKBP15* and *PUM1* expression. All data are the mean ± SD of three biological replicates, with significance determined by unpaired T-test. **E**, Western blot of TRIB1 and MYC from induced monoclonal cell lines over 96 hours. Gel was loaded with 50 μg total protein for each time point.

*TRIB1* knockdown, by siRNA SMARTpool transfection, in MDA-MB-231 (Supplementary Fig. 2C) and HCC1419 (Supplementary Fig. 2B) cells also significantly suppressed cell proliferation. In HCC1419 cells, which have the highest *TRIB1* copy number and expression levels, *TRIB1* knockdown resulted in a 1.6-fold (*P* = 0.0002, paired T-test) (Supplementary Fig. 4A-D) decrease in cell proliferation, with a 1.2-fold (*P* = 0.0256, paired T-test) decrease in cell number 120 hours after transfection (Supplementary Fig. 6D). Similarly, *TRIB1* knockdown in MDA-MB-231 cells, which are diploid at the *TRIB1* locus, resulted in a 1.6-fold (*P* = 0.0316, unpaired T-test) decrease in cell proliferation rate (Supplementary Fig. 5A-C) and a 1.3-fold (*P =* 0.0363, paired T-test) decrease in cell number 72 hours after transfection (Supplementary Fig. 6B). The consistent phenotype associated with *TRIB1* knockdown across cell lines with varying *TRIB1* copy number, suggests *TRIB1* is involved in breast cancer cell proliferation, independent of its amplification status.

Given the role of *MYC* in cell proliferation and the previously observed synthetic lethal relationship between *TRIB1* and *MYC* (20), the impact of *TRIB1* on *MYC* expression was examined in MCF7 cells. MCF7 cells carry *TRIB1/MYC* co-amplification, with an estimated five copies of each gene (Fig 1B). *TRIB1* knockdown resulted in a 2.1-fold (*P* = 0.0151) decrease in *MYC* expression, 72 hours after initial transfection (Fig. 2D). *TRIB1* knockdown by shRNA resulted in a sustained suppression of both TRIB1 and MYC protein levels (Fig. 2E) over a 96-hour period. TRIB1 and MYC protein levels increased over time when not suppressed by TRIB1 knockdown, suggesting both are important for the maintenance and promotion of MCF7 cell proliferation. However, *MYC* expression is known to be linked to changes in proliferation (37), therefore the mechanism linking *MYC* expression to *TRIB1* expression requires further investigation.

### qPLEX-RIME confirms TRIB1 associations with ubiquitination machinery and reveals associations with co-factors of key oncogenic transcription factors estrogen receptor and β-catenin

Quantitative Rapid Immunoprecipitation Mass Spectrometry of Endogenous proteins (qPLEX-RIME) (38) was used to determine the endogenous interactome of TRIB1 in breast cancer cells. This approach allows for the identification of key physical associations, without the bias of exogenous protein overexpression, or the need for introduction of exogenous tags. The TRIB1 qPLEX-RIME has the potential to both confirm TRIB1’s known physical associations, and reveal previously unreported interactions, providing insight into currently undescribed functions of TRIB1. Given the relatively high levels of TRIB1 observed in MCF7 and HCC1419 (Fig. 1D), qPLEX-RIME was conducted on these cell lines. The qPLEX-RIME workflow is illustrated in Fig. 3A: for each cell line four biological replicates were collected, and samples were labelled with TMT tags (39) to quantify relative enrichment of proteins compared to the IgG isotype control immunoprecipitations. 558 and 367 proteins were found to be significantly associated with TRIB1 in MCF7 and HCC1419 cells respectively (Log2FC > 2, (FDR) adj.*p-value* < 0.01). These associated factors were further refined by prioritizing those known to be localized to the nucleus, where TRIB1 has previously been reported to be located (7). This final filter resulted in 422 and 260 TRIB1- associated factors of interest from the MCF7 and HCC1419 cells. Comparison of the MCF7 and HCC1419 qPLEX-RIME datasets revealed 105 TRIB1-associated proteins common between the two cell lines, which were grouped by function based on Uniprot function description (Fig. 3B).

**Figure 3:**
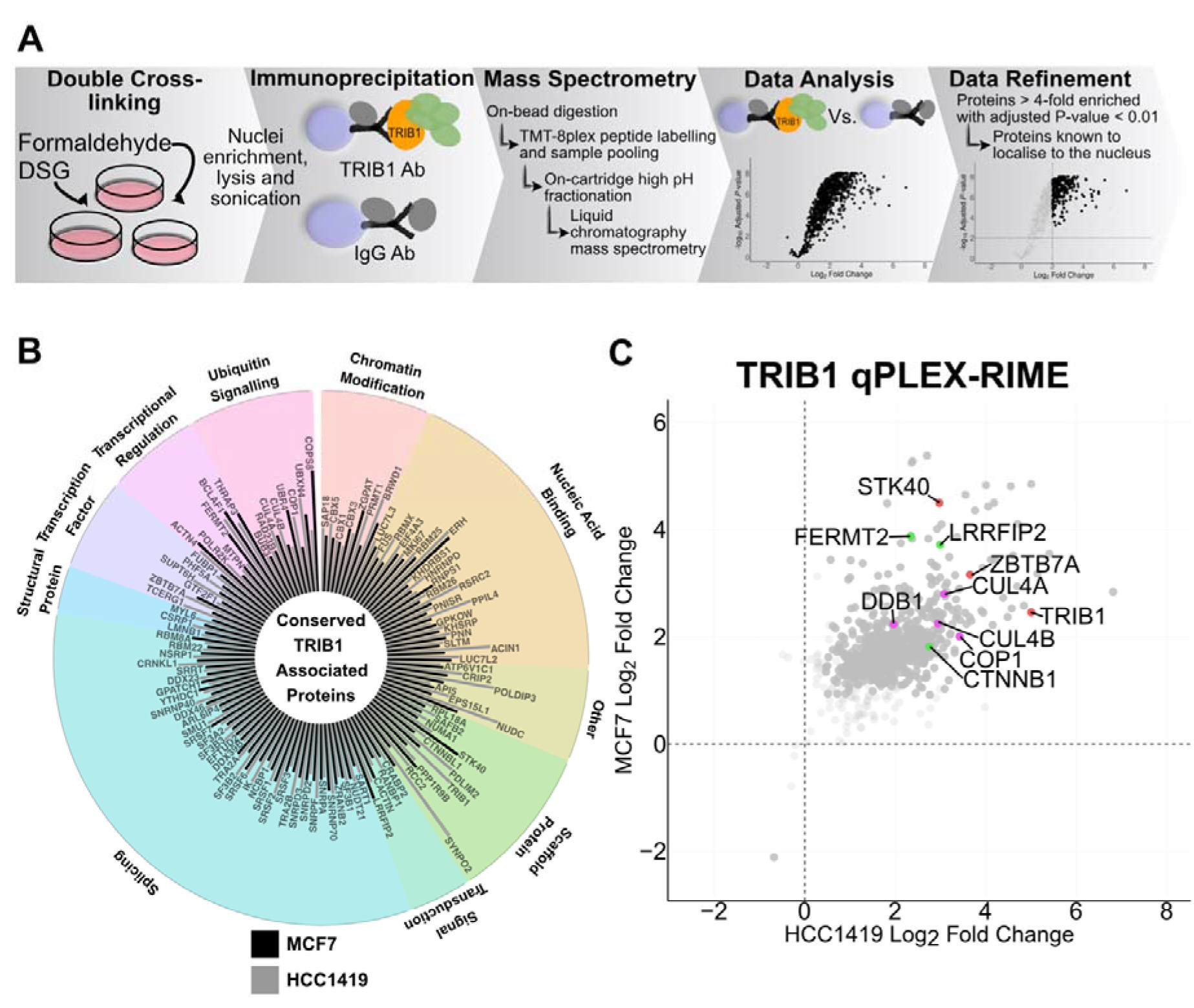
qPLEX-RIME of TRIB1 in MCF7 and HCC1419 cells provides the first endogenous interactome of TRIB1 in breast cancer cells. **A**, Schematic showing the workflow for TRIB1 qPLEX-RIME. **B**, Summary of the significantly enriched (Log2FC *> 2,* adj*.p-value* < 0.01) TRIB1 associated factors conserved in both the MCF7 and HCC1419 datasets. Bar height is representative of log_2_ fold-enrichment. **C**, Comparison of all factors identified in both the MCF7 and HCC1419 qPLEX-RIME datasets. Light grey are non-significant (Adj.*p-value* > 0.01), dark grey are significant (Adj.*p-value* < 0.01), purple are ubiquitination related factors, green are WNT/β-catenin related factors, orange are other factors of interest.

Consistent with the well-characterised association between TRIB1, COP1 and CUL4- associated ubiquitin ligases in other cell-types (5, 7, 40, 41), we observed COP1 as a prominently enriched TRIB1-partner, along with CUL4(A and B), and DDB1 (Fig. 3B). This suggests that the degradative role of TRIB1 may be at least partially conserved in breast cancer cells, as TRIB1 associates with the same suite of ubiquitin-ligase partners.

As TRIB1 regulates transcription via the ubiquitination of transcription factors in myeloid cells and other contexts, we focused on investigating its link to transcriptional regulation in breast cancer. Examination of TRIB1-associated transcription factors and transcriptional regulators revealed two factors of particular interest, the β-catenin co-factor FERMT2 (MCF7 log_2_FC = 3.88, adj.*p-value* < 0.01 HCC1419 log_2_FC = 2.34, adj.*p-value* < 0.01) (42, 43), and the transcription factor ZBTB7A (MCF7 log_2_FC = 3.16 adj. *p-value* < 0.01), HCC1419 log_2_FC = 3.65, adj.*p-value* < 0.01) (Fig. 3B). In addition, TRIB1 associated with β-catenin, and the upstream WNT signal enhancer LRRFIP2 (MCF7 log_2_FC = 3.72, adj. *p-value* < 0.01 HCC1419 log_2_FC = 2.99, adj. *p-value* < 0.01) (Fig. 3B, C, Supplementary Table 1). qPLEX-RIME also revealed associations with a selection of chromatin modifiers, including enzymatically active proteins such as the methyltransferase PRMT1, and the heterochromatin proteins CBX1, CBX3 and CBX5 (Fig. 3B, Supplementary Table 1).

### Transcriptional profiling implicates TRIB1 in cell-cycle regulation

As our qPLEX-RIME analysis indicated that TRIB1 associates with a range of potent transcriptional regulators, we sought to assess the transcriptional changes induced by reducing *TRIB1* expression. RNAseq was performed on two MCF7 *TRIB1* knockdown models. RNAseq of *TRIB1* knockdown by siRNA SMARTpool transfection (Supplementary Fig. 8A), and inducible stably transduced single shRNA knockdown (Supplementary Fig. 8B) were analyzed independently for differentially expressed genes. To capture the early changes driven by *TRIB1* knockdown, rather than the longer-term consequences of altered *MYC* expression (Fig. 2D), RNA was collected 48 hours after siRNA transfection or shRNA induction. *TRIB1* was significantly downregulated at 48 hours in both siRNA (*P <* 0.0001, unpaired T-test) and shRNA (*P* = 0.0019, unpaired T-test) models (Supplementary Fig. 8), while *MYC* expression was not significantly changed (Supplementary Fig. 9B).

To identify cellular processes altered by *TRIB1* knockdown, Fast Gene Set Enrichment Analysis (FGSEA) (44) was performed using the Hallmark signature collection (45). In total, 23 of the 50 Hallmark signatures were significantly enriched (FDR adj.*p-value* < 0.2, exploratory *P*-value threshold) in both the siRNA and shRNA knockdown gene sets (Fig. 4A). Consistent with the previously observed phenotype of reduced proliferation, signatures positively associated with proliferation (“G2M Checkpoint”, “E2F Targets”, “MYC Targets V1” and “MYC Targets V2”) were significantly enriched with genes downregulated by both siRNA and shRNA *TRIB1* knockdown (Fig. 4A). Interestingly, the “p53 Pathway” signature, which is negatively associated with proliferation, was upregulated by *TRIB1* knockdown (Fig. 4A). These results suggest cell cycle regulation as the mechanism behind the proliferation inhibition phenotype observed (Fig. 2A, B, C).

**Figure 4:**
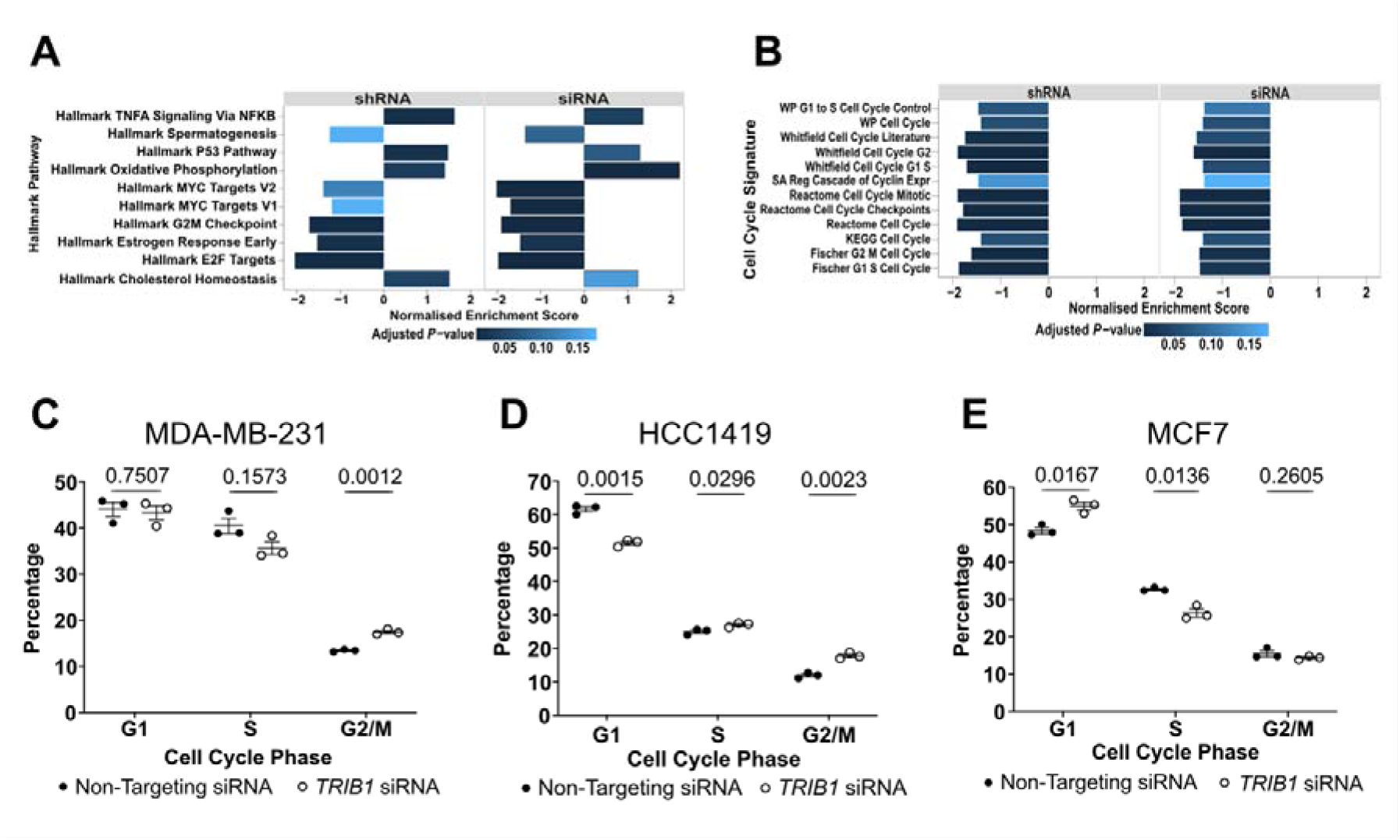
*TRIB1* knockdown causes expression changes in MCF7 cells, and inhibits cell cycle progression in MCF7, MDA-MB-231 and HCC1419 cells. RNAseq was performed on MCF7 cells after 48 hours of *TRIB1* knockdown by siRNA SMARTpool transfection, or shRNA induction with 2 μg/mL for monoclones (GSE233138). Differential expression analysis was performed independently on the siRNA and shRNA datasets. The resulting genesets were ordered by T-statistic for Fast GeneSet Enichment Analysis (FGSEA). **A**, Comparison of significantly enriched Hallmark pathways from FGSEA with MCF7 siRNA and shRNA *TRIB1* knockdown genesets. Significance threshold set at FDR adj.*p-value* < 0.2. **B**, FGSEA of cell cycle related signatures using the MCF7 siRNA and shRNA *TRIB1* knockdown genesets. Significance threshold set at FDR adjusted *P-*value < 0.2. Cell cycle analysis, 48 hours after transfection with the *TRIB1* or Non-Targeting ON-TARGETplus siRNA SMARTpools, in **C**, MCF7. **D**, MDA-MB-231 and **E**, HCC1419 cells. Cell cycle phase determined by propidium iodide staining with FACS data collection with the Guava benchtop flow cytometer. Cell cycle analysis performed using FlowJo software, data are the mean ± SEM of three biological replicates, each the mean of three technical replicates per treatment. Significance determined using multiple unpaired T-tests with FDR multiple testing correction.

The “G2M Checkpoint” and “E2F Targets” signatures were two of the most significantly enriched Hallmark signatures with strongly negative normalized enrichment scores in both gene sets (siRNA adj. *p-value* = 1.38e-3, shRNA adj. *p-value* = 4.35e-3) (Fig. 4A, Supplementary Table 2). To further validate these changes, targeted cell cycle related signatures were obtained from the Molecular Signature Database (MSigDB) and analyzed using FGSEA. Signatures of both G_2_-M, and G_1_-S phase transition were enriched with genes down-regulated by *TRIB1* knockdown (Fig. 4B). The downregulation of genes in the broad KEGG and Reactome cell cycle signatures further implicates *TRIB1* as a cell cycle regulator in breast cancer cells (Fig. 4B). The effect of *TRIB1* knockdown on cell cycle progression was examined using DNA content assessment by flow cytometry in MCF7, HCC1419 and MDA-MB-231 cells treated with siRNA SMARTpools.

### *TRIB1* knockdown induces cell cycle inhibition

*TRIB1* has previously been implicated as a regulator of G_1_-S phase transition in MDA-MB-231 cells released from MEK1/2 inhibition-induced cell cycle arrest (11). Consistent with that observation, flow cytometry analysis of cycling, unsynchronized, MDA-MB-231 cells revealed cell cycle inhibition in response to *TRIB1* knockdown (Fig. 4C-E). However, significant enrichment of cells in the G_2_/M phase of the cell cycle rather than G_1_-phase was observed (Fig. 4C). Cell cycle inhibition in response to *TRIB1* knockdown was also observed in HCC1419 cells with significant enrichment of cells in the G_2_/M phase (Fig. 4E). In addition, significant depletion of the cells in the G_1_ phase was observed in the TRIB1 knockdown population (Fig. 4E). In contrast, in MCF7 cells *TRIB1* knockdown led to a significant enrichment of cells in the G_1_ phase of the cell cycle, and a trend towards depletion of cells in S-phase suggesting inhibition at the G_1_-S phase transition (Fig. 4D).

While the association of *MYC* amplification with aggressive tumors is well described, evidence that *TRIB1* also drives cell cycle progression across breast subtypes further implicates *TRIB1* amplification as an essential component of 8q24 amplification, rather than simply a passenger of *MYC* amplification.

### TRIB1 knockdown is associated with decreased β-catenin, increased FERMT2 and a reduction in β-catenin associated transcription

In assessing potential mechanisms underlying TRIB1 cell-cycle regulation in breast cancer cells, we noted that TRIB1 associated with the key oncogenic transcription factor β-catenin, and the β-catenin co-factor FERMT2 (Fig. 4B, C). In addition, the Hallmark signature “WNT Beta Catenin Signaling” was significantly enriched with downregulated genes in the siRNA *TRIB1* knockdown gene set (Supplementary Table 2).

To further explore this link, we investigated the effect of *TRIB1* knockdown on β-catenin transcription using a targeted set of WNT/β-catenin related signatures from the MSigDB. Signatures related to WNT/β-catenin activity including the “Reactome Signaling by WNT” and “KEGG WNT Signaling Pathway” were significantly enriched with genes downregulated in the siRNA *TRIB1* knockdown gene set (Fig. 5A). Many of these pathways also had negative enrichment scores from FGSEA with the shRNA *TRIB1* knockdown gene set (Fig 5A). To confirm the regulation of β-catenin driven transcription, targeted assessment of β-catenin targets *AXIN2* and *FSCN1* was performed using RT-qPCR. Both showed significant downregulation in MCF7 cells – *AXIN2* expression decreased by 2.6-fold (*P =* 0.0007) and 1.7-fold (*P =* 0.0186) upon siRNA and shRNA *TRIB1* knockdown respectively. Similarly, *FSCN1* expression decreased by 2.5-fold (*P =* 0.0069) or 1.8-fold (*P =* 0.132) in response to siRNA or shRNA *TRIB1* knockdown (Fig. 5B, C).

**Figure 5:**
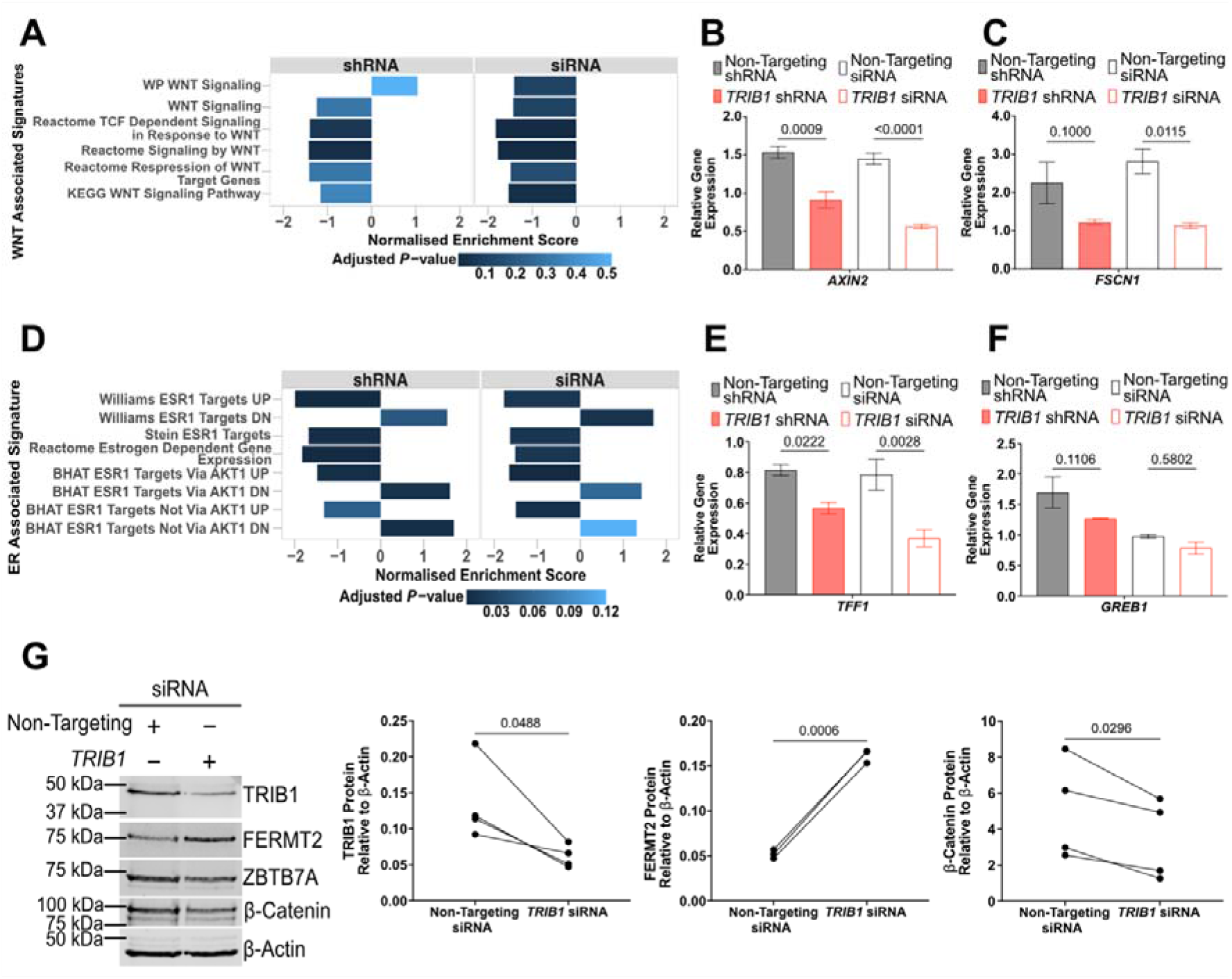
*TRIB1* knockdown suppresses ER and β-catenin driven transcription in MCF7 cells. **A**, FGSEA of ER related signatures using the MCF7 siRNA and shRNA *TRIB1* knockdown genesets. Significance threshold set at FDR adjusted *P-*value < 0.2. **B**, *TFF1* and **C**, *GREB1* expression, in MCF7 cells, was determined by RT-qPCR after 48 hours of *TRIB1* knockdown by siRNA SMARTpool, or shRNA induction. Same RNA as was used for the RNAseq analysis. Expression was normalized to the geometric mean of *PUM1* and *FKBP15* expression. All data are the mean ± SEM of three biological replicates, with the significance determined using multiple unpaired T-tests with FDR multiple testing correction. **D**, FGSEA of WNT/β-catenin related signatures using the MCF7 siRNA and shRNA *TRIB1* knockdown genesets. **E**, *AXIN2* and **F**, *FSCN1* expression, in MCF7 cells, was determined by RT-qPCR after 48 hours of *TRIB1* knockdown by siRNA SMARTpool, or shRNA induction. Same RNA as was used for the RNAseq analysis. Expression was normalized to the geometric mean of *PUM1* and *FKBP15* expression. All data are the mean ± SEM of three biological replicates, with the significance determined using multiple unpaired T-tests with FDR multiple testing correction. **G**, Representative western blot of TRIB1, ZBTB7A, FERMT2 and β-catenin from MCF7 cells 72 hours after transfection with the *TRIB1* or Non-Targeting ON-TARGETplus siRNA SMARTpools. Densitometry for TRIB1, FERMT2 and β-catenin was normalized to β-actin densitometry. Data points are individual biological replicates, with TRIB1 and non-targeting siRNA treated lysate from the same replicate run on the same gel. Significance determined by paired T-test.

*TRIB1* knockdown in MCF7 cells resulted in significantly (*P* = 0.0006) increased FERMT2 protein levels (Fig. 5G). Total β-catenin levels were also assessed and were decreased (*P* = 0.0296) in response to *TRIB1* knockdown (Fig. 5G). These results suggest TRIB1 may influence the stoichiometry, and hence activity and stability of the β-catenin transcriptional complex.

### TRIB1 regulates estrogen receptor driven transcription contributing to the modulation of the cell cycle in MCF7 cells

As an ER+ cell line, the primary driver of MCF7 cell proliferation is the estrogen receptor (46). A physical association between TRIB1, and the known ER co-factor ZBTB7A (38, 47) observed in qPLEX-RIME (Fig. 3B, C) providing a potential mechanism through which TRIB1 could influence ER-associated transcription, and MCF7 cell proliferation. In contrast to FERMT2, no significant change in ZBTB7A protein level was observed in response to *TRIB1* knockdown in MCF7 cells (Supplementary Fig. 13A-D). However, despite the lack of protein level changes in ZBTB7A genes in the estrogen response, Hallmark signatures tended to be downregulated in response to *TRIB1* knockdown, suggesting TRIB1 may contribute to estrogen-dependent gene expression in MCF7 cells (Fig. 4A, Supplementary Table 2). This suggests that – in contrast to many TRIB1 associated proteins described previously (5, 8, 9, 26, 48–50) – TRIB1 does not appear to mediate targeted degradation of ZBTB7A in MCF7 cells, but likely influences ER driven transcription through ZBTB7A via an alternative mechanism.

Further exploration of ER related expression using a range of signatures from the MsigDB revealed modulation of ER-associated activation and repression (Fig. 5D). Signatures of ER repressed genes were enriched with upregulated genes in both *TRIB1* knockdown gene-sets, while signatures of ER activated genes tended to be downregulated (Fig. 5D). Suppression of ER target genes was confirmed by RT-qPCR, with 2.1- or 1.4-fold decrease in expression of the key estrogen responsive gene *TFF1* in response to siRNA or shRNA *TRIB1* knockdown respectively (Fig. 5E, F). Expression of *GREB1* also decreased in response to *TRIB1* knockdown, although this change was not statistically significant (Fig. 5E, F). These results were reproduced in biologically independent experiments using siRNA *TRIB1* knockdown, validating the observed effect of *TRIB1* knockdown on ER driven transcription (Supplementary Fig. 9E, F). This modulation of ER driven transcription likely has a strong influence over MCF7 cell proliferation, contributing to the *TRIB1* knockdown phenotype observed (Fig. 2A-C).

## Discussion

The association of *TRIB1/MYC* amplification with poor patient outcome (Fig. 1A) (31) highlights the need to understand the role of *TRIB1* in breast cancer. Co-amplification of *TRIB1* with *MYC* was observed in three different breast cancer cell lines (Fig. 1B), with *TRIB1* expression and protein levels being proportional to copy number in the cell lines investigated (Fig. 1C, D). *TRIB1* knockdown inhibited cell cycle progression, slowing cell proliferation in MCF7, HCC1419 and MDA-MB-231 breast cancer cells (Fig. 2A-C, Fig. 4C-E, Supplementary Fig. 3-6), indicating the role of *TRIB1* in breast cancer cell proliferation is at least partially independent of its amplification status. Examination of the TRIB1 interactome using qPLEX-RIME in HCC1419 and MCF7 cells (Fig. 3) highlighted the conservation of the well-characterised physical association between TRIB1 and COP1 (5, 7, 9, 51), and revealed an association with the wider CUL4 ubiquitination machinery (Fig. 3B). A number of physical associations with key regulatory transcription factors, including core components of β-catenin transcriptional complex, were also identified in the TRIB1 interactome. In line with the interactome, *TRIB1* knockdown influenced cell cycle related, ER driven, and β-catenin driven transcription in MCF7 cells (Fig. 4A-B, 5A-F). The assessment of β-catenin related TRIB1 associated factors after *TRIB1* knockdown suggests that TRIB1 is likely to regulate β-catenin transcription by facilitating degradation of the β-catenin co-factor FERMT2 (Fig. 5G). Levels of the ER co-factor ZBTB7A were not affected by *TRIB1* knockdown in MCF7 cells (Fig. 5G), potentially suggesting a novel mechanism of TRIB1 in the regulation of ER-driven transcription, although further investigation is required.

*TRIB1* knockdown inhibits breast cancer proliferation and *MYC* expression in MCF7 cells, where *TRIB1* and *MYC* are co-amplified. These results suggest *TRIB1* amplification may support amplification-driven *MYC* overexpression, a function which would make *TRIB1* an appealing target, not only in breast cancer, but across a range of cancer types. Downregulation of β-catenin and ER transcriptional programs provide two possible avenues for the regulation of *MYC* expression, and implicates TRIB1 as critical in the maintenance of oncogenic transcriptional programs.

The TRIB1-COP1 association was originally identified in leukemia (1, 5–9), and is central to TRIB1’s ability to mediate degradation is conserved in breast cancer cells, suggesting conservation of the COP1 dependent transcription factors degradation mechanism of TRIB1 transcriptional regulation observed in AML (3, 5, 6, 52). Examination of the wider TRIB1 interactome showed association with the CUL4A/B ubiquitination machinery (Fig. 3B), implicating TRIB1 as a key adaptor protein for CUL4 complex, which is a central ubiquitin dependent degradation mediator. Given the structural similarity between TRIB1, TRIB2 and STK40, the potential for the context-specific association of these proteins with the CUL4A/B complex should be further investigated.

Consistent with the role of TRIB1 as a regulator of transcription factor abundance in haemopoietic differentiation and dysregulation (3, 5, 6, 61), as well as the strong association with ubiquitination machinery, TRIB1 was found to associate with a selection of transcription factors and co-factors (Fig. 3B). Among these associations, of particular interest were the associations with β-catenin and FERMT2, known members of the β-catenin transcriptional complex, which is a key driver of *MYC* expression (19, 53–55). FERMT2 protein levels increased after *TRIB1* knockdown (Fig 5G), suggesting that TRIB1 may be involved in the degradation FERMT2. Given the conservation of the required associations with ubiquitination machinery, TRIB1 likely mediates FERMT2 degradation through its canonical scaffolding mechanism of mediating an interaction between a target protein and COP1 (56). While TRIB1-mediated FERMT2 degradation is apparent (Fig. 5G), how TRIB1 affects β-catenin complex stoichiometry and stability remains unclear. Given the apparent destabilization of β-catenin upon TRIB1 knockdown (Fig. 5G), it is possible that TRIB1 could interact independently with β-catenin or FERMT2, or may scaffold the two proteins. Regulation of FERMT2 and β-catenin also provides a potential mechanism through which TRIB1 could act in metabolic signaling as TRIB1, FERMT2 and β-catenin dependent WNT signaling are all implicated in lipid regulation (26, 57–62).

The regulation of β-catenin-driven transcription provides a mechanism for the regulation of *MYC* expression across genetic backgrounds. However, in ER+ MCF7 cells, one of the most potentially impactful effects of *TRIB1* knockdown observed was the suppression of ER- mediated transcription and repression (Fig. 5A). ER is known to bind upstream of *MYC* and a range of cell cycle related genes including *MYB, CCND1* and *BCL2* (63). Therefore, in MCF7 cells, regulation of ER is a potential mechanism for both the cell cycle inhibition, and *MYC* suppression observed upon TRIB1 knockdown (Fig. 2D, 4E). The known ER co-factor, and chromatin remodeler ZBTB7A, was found to be associated with TRIB1 (Fig. 3B) but no change in ZBTB7A levels was observed in response to *TRIB1* knockdown (Fig. 5G, Supplementary Fig. 13A-D). Consequently, it seems likely that that TRIB1 regulation of ER signaling occurs independently of COP1. ZBTB7A is an important regulator of promoter accessibility for a range of transcription factors, including ER (47, 64–67). To regulate chromatin accessibility, ZBTB7A co-operates with a range of chromatin remodelers and epigenetic modifiers (64, 66–68). Further TRIB1-associated epigenetic modifiers include the heterochromatin protein family (CBX1/3/5), the methyltransferase PRMT1, and the SIN3-repressor complex subunit, SAP18 (Fig. 3B). The association with both a localizing transcription factor and key effector proteins, suggests TRIB1 may facilitate the formation of chromatin modifying complexes at enhancer and promoter sites, by scaffolding their association with ZBTB7A. This potential relationship between ZBTB7A and TRIB1, may allow TRIB1 to influence a range of different transcriptional programs across a variety of different genomic backgrounds, including both ER- and ER+ breast cancers (1, 2, 69).

This study demonstrates that TRIB1 is involved in the regulation of cell proliferation and the cell cycle across breast cancer subtypes (9). This first interrogation of the endogenous TRIB1 interactome in breast cancer cells, coupled with global assessment of the transcriptional consequences of *TRIB1* knockdown, have revealed two major mechanisms through which TRIB1 may regulate cell proliferation. Further investigation of TRIB1 in β-catenin transcription and ZBTB7A function is critical for understanding the co-operative role of TRIB1 with MYC in breast cancer and more broadly.

## Materials and Methods

### Cell Culture

The MDA-MB-231 (Cat# HTB-26), MCF7 (Cat# HTB-22) and HCC1419 (Cat# CCRL-2326) breast cancer cell lines were obtained from ATCC. MCF7 and MDA-MB-231 cells were cultured in RPMI 1640 media (Sigma-Aldrich) with 10% fetal bovine serum (FBS, Gibco). HCC1419 cells were cultured in RPMI 1640 Hybi-Max media (Sigma-Aldrich) with 10% FBS. Media with FBS is termed ‘full media’. All cell lines were grown in a humidified 37°C incubator with 5% carbon dioxide.

### RNA collection and RT-qPCR

Total RNA was extracted from cell lines using the Zymo Quick RNA Mini-Prep kit, or the RNAGEM purification kit according to the manufacturer’s instructions. RNA quality and quantity were assessed using a NanoDrop ND-1000 and Qubit Fluorometer (both ThermoFisher). RNA was reverse transcribed using the PrimeScript RT Reagent Kit (Takara). RT-qPCR was performed with TaqMan Gene Expression Master Mix (Thermo Fisher) using predesigned PrimeTime qPCR Probe Assays for FKBP15 (Hs.PT.49a.2552313), PUM1 (Hs.PT.56a.1997572), TRIB1 (Hs.PT.58.2667458), MYC (Hs.PT.58.26770695), AXIN2 (Hs.PT.58.39305692), FSCN1 (Hs.PT.58.614427), GREB1 (Hs.PT.58.26216464) and TFF1 (Hs.PT.58.168461) from IDT. Gene expression of the target genes was normalized to the geometric mean of the reference genes FKBP15 and PUM1.

### siRNA Transfection

Cells were seeded at an appropriate density 24 hours prior to transfection. Cells were transfected with the *TRIB1* targeting (L-003633-00-0010) or non-targeting control (D- 001810-10-2) ON-TARGETplus siRNA SMARTpools (Dharmacon) using RNAiMAX (Thermo Fisher) according to the manufacturer’s instructions. Transfection mixes were made in phenol red free RPMI 1640 base media and added to cells in a dropwise manner to the cells in full media. Cells were incubated with the transfection mixture for 24 hours before it was removed by way of a full media change.

### Antibodies

**Table of primary antibodies used for Western Blot. All antibodies were diluted in TBST + BSA (5% w/v).**

| Antigen | Catalogue number | Supplier | Host Species | Concentration |  |
| --- | --- | --- | --- | --- | --- |
| TRIB1 | 8052 | Genentech | Rat | 1:2000 |  |
| MYC | 9402 | Cell Signaling Technology | Rabbit | 1:1000 |  |
| β-catenin | 9562 | Cell Signaling Technology | Rabbit | 1:1000 | |
| FERMT2 | OTI9E4 | Invitrogen | Mouse | 1:2000 |  |
| ZBTB7A | ab175918 | Abcam | Rabbit | 1:5000 |  |
| α-tubulin | 05-829 | Millipore | Mouse | 1:5000 | |
| β-actin | A2066 | Sigma-Aldrich | Rabbit | 1:2000 | |

The secondary antibodies were rabbit anti-rat HRP-conjugate (Thermo Fisher) used at 1:5000 in TBST + BSA (1% w/v), and anti-mouse 800CW (926-32210) and anti-rabbit 680RD fluorescent secondaries (LI-COR) diluted between 1:7500 to 1:10000 in TBST + BSA (1% w/v).

The antibodies TRIB1 (PA5-66650) and IgG control (ab171870) used for qPLEX-RIME immunoprecipitations, were from Thermo Fisher and Abcam respectively.

### Western Blot Analysis

Total protein was collected from cell lines using an EasyPep lysis buffer mix (per 100 μL, 98.5 μL EasyPep Lysis Buffer (ThermoFisher), 1 μL 100X EDTA-free Protease Inhibitor (Roche), 0.5 μL Pierce Universal Nuclease (ThermoFisher)). Total protein concentration was determined using the Pierce BCA assay kit (ThermoFisher). The desired amount of protein was mixed with SDS sample buffer and run on 12-18% gradient SDS-PAGE gel at 220V for 55 minutes in a water-cooled tank. Proteins were transferred onto a nitrocellulose membrane (ThermoFisher) using the iBlot 2 dry transfer system (ThermoFisher) with a multi-step transfer protocol (one minute at 20V, four minutes at 23V, one minute at 25V). After transfer, the membrane was incubated at 4°C for one hour in blocking buffer (Tris-buffered saline with 0.1% Tween 20 (TBST), 5% BSA (w/v)). Primary antibody incubations were performed overnight at 4°C. Following incubation, blots were washed three time, for five minutes, in TBST. All secondary antibody incubations were performed for one hour, at 4°C then the blot washed as above before imaging. The LI- COR Odyssey FC imaging system (LI-COR Biosystems) was used for imaging. Fluorophore conjugated secondary antibodies were imaged directly by two-minute exposure in the relevant fluorescence imaging channel. Where a horseradish peroxidase (HRP) conjugated secondary was used, it was developed using Pierce ECL Western Blotting Substrate (ThermoFisher) according to the manufacturer’s instructions and imaged by 10-minute exposure in the chemiluminescence channel. Densitometry was performed using Image Studio Lite software (LI-COR Biosystems), and all densitometry was normalized to β-actin or ⍺-Tublin. Where all densitometry was collected from a single blot one-way ANOVA was used to test statistical significance with each repeat on the blot being a biological replicate. Where biological replicates (*TRIB1* knockdown experiments) where spread across multiple western blots significance was determined by paired T-test with pairing of each individual biological replicate. Unless stated otherwise all biological replicates consisted of a single lane per condition. Statistical tests were performed using Prism 9.

### IncuCyte Live Cell Imaging

Cells were seeded in full growth media, in 96-well plates, at the cell line specific density (MCF7, 1000 cells; HCC1419, 15000 cells; MDA-MB-231, 2000 cells), and allowed to adhere overnight. Cells were transfected in triplicate with 1 pmol of the relevant siRNA SMARTpool (Dharmacon), using 0.2 μL of RNAiMAX (ThermoFisher) per well according to the manufacturer’s instructions, immediately placed in the IncuCyte live cell imager at 37◦C, 5% CO_2_ and a protocol of two hourly imaging was started. The media was changed 24 hours after transfection, then every 72 hours thereafter and imaging continued for a minimum of 168 hours. Average cell confluency and standard deviation was calculated from images every two hours using IncuCyte software. Confluency over 168 hours was plotted using Prism 9 software (GraphPad). Confluency change per hour was calculated based on the linear growth phase for each condition and plotted using Prism 9. Significance of difference in change in confluency per hour was determined by paired T-test when three or more biological replicates were performed. Samples were paired based on biological replicate (each repeat was an independent passage and transfection). Where only two biological replicates were available significance was determined by unpaired T-test. Statistical tests were performed using Prism 9.

### Nuclei Staining and Counting

Cells were seeded in 96-well plates, at cell line specific density (MCF7, 1000 cells; HCC1419, 15000 cells; MDA-MB-231, 2000 cells), and allowed to adhere overnight. Cells were transfected as described above, with reagent controls in duplicate at the same concentration. Media was changed after 24 hours, and MCF7 and MDA-MB-231 cells were incubated for a further 48 hours before being fixed and stained with 1 μg/mL Hoechst, 0.25% paraformaldehyde and 0.075% saponin in PBS. The HCC1419 cells were incubated for 72-hours post media change before fixing and staining. Stained cells were imaged using the Cytation 5 (Biotek) at 4X magnification taking six fields per well. Nuclei were counted using Gen5 software (Biotek) and normalized to the siRNA and RPMI reagent control nuclear counts. Significance was determined by unpaired T-test where technical replicates were averaged for each biological replicate and each cell line was test independently.

For nuclei counting of MCF7 monoclones with inducible *TRIB1* targeted and control shRNA, cells were treated with 2 μg/mL doxycycline in complete media for 72 hours prior to seeding to induce shRNA expression. Cells were seeded at 1000 cells/well in media containing 2 μg/mL doxycycline and grown for 120 hours. Every 24 hours, six wells for each shRNA were stained with 1 μg/mL Hoechst in PBS for 30 minutes and imaged using the Cytation 5 (Biotek) at 4X magnification taking six fields per well. Nuclei were counted using Gen5 software (Biotek). Significance of the difference in nuclei count at each time point was determined by multiple unpaired T-tests on three biological replicates, with Holm-Šídák multiple testing *P-value* correction. Six technical replicates were averaged for each biological replicate. Statistical tests were performed using Prism 9.

### Double cross-linking for qPLEX-RIME

Cells were seeded in 10 cm dishes and allowed to grow for 72 hours. For crosslinking, media was removed and cells incubated with 20 mL of PBS containing 2 mM disuccinimidyl glutarate (DSG) (Santa Cruz Chemical) for 20 minutes at room temperature. DSG was removed, and cells incubated with 10 mL of 1% methanol free formaldehyde (ThermoFisher) in PBS for 10 minutes. The cross-linking reaction was quenched by adding glycine to a final concentration of 0.125 M. Cells were washed twice with ice cold PBS then scraped in 500 μL of PBS + protease inhibitors (PIs) (Roche). Cells were centrifuged at 8000 rpm (4^◦^C) for three minutes, washed again in PBS + PI, and supernatant removed. Cell pellets were snap frozen on dry-ice and stored at −80^◦^C.

### Nuclear Fractionation and Immunoprecipitation for qPLEX-RIME

Twenty-four hours prior to cell lysis, 10 μg of the respective antibodies were bound to 100 μL of Protein A magnetic beads (ThermoFisher). Beads were blocked with three 1 mL washes of 5 mg/mL BSA in PBS. After blocking, beads were suspended in 500 μL of PBS + BSA and 10 μg antibody added. Beads were rotated at 20 rpm at 4^◦^C overnight.

For the exaction of the nuclear protein fraction, cell pellets were thawed on ice, then resuspended in Lysis Buffer (LB1) (50 mM HEPES-KOH (pH 7.5), 140 mM NaCl, 1 mM EDTA, 10% glycerol, 0.5% NP-40/Igepal CA-630, 0.25% Triton X-100) supplemented with protease inhibitors (Roche) (1 mL per 15 cm dish), and rotation mixed at 4^◦^C for 10 minutes. Nuclei were pelleted by 2000 RCF centrifugation at 4^◦^C for five minutes. The supernatant was removed, the pellet resuspended in LB2 (10 mM Tris-HCl (pH 8.0), 200 mM NaCl, 1 mM EDTA, 0.5 mM EGTA) supplemented with protease inhibitor (1 mL per 15 cm dish) and rotation mixed at 4^◦^C for five minutes. Nuclei were pelleted as before and resuspended in LB3 (10 mM Tris-HCl (pH 8.0), 100 mM NaCl, 1 mM EDTA, 0.5 mM EGTA, 0.1% sodium deoxycholate, 0.5% sodium lauroylsarcosine) supplemented with protease inhibitors (300 μL per 15 cm dish) and sonicated at 4^◦^C (Biorupter by Diagnode) for 15 cycles of 30 seconds on and 30 seconds off, or until chromatin was fragmented down to 200-300 bp. After fragmentation, Triton X-100 was added to a final concentration of 1% and the solution was centrifuged at 20 000 rcf for 10 minutes at 4^◦^C. During centrifugation, antibody beads were washed three times with PBS + BSA (5 mg/mL) and then resuspended in 200 μL of LB3 with 1% Triton X-100 and PI. Supernatant was transferred to a fresh 2 mL tube, beads added and incubated at 4^◦^C overnight on a mixing rotor.

For qPLEX-RIME experiments, four 15 cm dishes of cells were used per antibody per biological replicate. Biological replicates were pooled in eight 15 cm dish batches during LB1 resuspension and then split equally across TRIB1 and IgG antibodies for immunoprecipitation.

After overnight immunoprecipitation, beads were washed on ice 10 times with 1 mL RIPA buffer [50 mM HEPES (pH 7.6), 1 mM EDTA, 0.7% sodium deoxycholate, 1% NP-40/Igepal CA-630, 0.5 M LiCl]. Then beads were washed twice in 500 μL of 100 mM ammonium bicarbonate (AMBIC), and transferred to a fresh tube during the second wash. AMBIC was removed and beads were snap frozen on dry-ice.

### Trypsin Digestion, TMT-labelling and LC-/MS analysis for qPLEX-RIME

On beads trypsin digestion, TMT-labelling and mass spectrometry analysis were carried out using the methods previously described by Papachristou et al, 2018 (38). Briefly, samples were digested with trypsin followed by a peptide clean-up with Ultra-Micro C18 Spin Columns (Harvard Apparatus). Peptides were dried and labelled with TMT-10plex reagents (Thermo Scientific) and purified using Reversed-Phase spin columns (Pierce #84868). Nine fractions were collected and run on a Dionex UltiMate 3000 UHPLC system coupled with the nano-ESI Fusion Lumos (Thermo Scientific). The 10 most intense fragments were selected for Synchronous Precursor Selection (SPS) HCD-MS3 analysis with MS2 isolation window 2.0 Th. The HCD collision energy was set at 55% or 65% and the detection was performed with Orbitrap resolution 50K and in scan range 100–400 m/z. Data was processed using the Proteome Discoverer 2.1 and the qPLEXanalyzer (38) R package was used for downstream statistical analysis. Differential enrichment between the TRIB1 and IgG pulldowns was determined using Limma with FDR multiple testing *P-value* adjustment.

### RNAseq Analysis

MCF7 cells or monoclones 6080023-3 and Non-Targeting-12 were seeded at 100 000 cells/well in six well plates and allowed to adhere overnight. MCF7 cells were transfected in triplicate and shRNA expression induction was induced in monoclones by treatment with 2 µg/mL doxycycline. RNA was collected 48 hours as described above.

Three biological replicates of each knockdown and matched control were prepared for transcriptomic analysis, and RNA submitted to Otago Genomics for cDNA conversion, library generation and sequencing. Sequence was provided by Otago Genomics via Illumina BaseSpace Sequencing Hub. Initial sequence quality was assessed by Otago Genomics using FastQC (70) and summarized with MultiQC (71) software. Sequence was aligned to the GRCh38 primary assembly (downloaded 23/05/2020) of the human genome using the STAR ultrafast universal RNAseq alignment software (72). The quality of the alignment was determined using FastQC (70) and summarized with MultiQC (71) software. Aligned sequences were imported into Rstudio (R version 3.6) (73, 74) and feature counting was performed with the NCBI RefSeq annotation of the human genome (build hg38.2) (75), using Rsubread (76, 77).

Feature counts were normalized and transformed using the voom analysis pipeline (78), and differential expression was performed independently on the siRNA and shRNA expression data sets using Limma (79, 80) in RStudio. Multiple testing correction for differential expression analysis was performed using FDR for *P-value* adjustment. Gene Set Enrichment Analysis (GSEA) of signature databases was performed using FGSEA (44) in RStudio, using FDR for multiple testing *P-value* correction.

### Cell Cycle Analysis

Cells were seeded and transfected 24 hours later as previously described. Cells were collected 48 hours after transfection and fixed with ice cold 70% ethanol. For DNA content based assessment of cell cycle, cells were permeabilized in PBS buffer (1 mM EDTA, 0.1% Triton X-100) and treated with RNase A (0.05 mg/mL) for 20 minutes at 37^◦^C, then incubated with PBS buffer + PI (1 µg/mL) for 15 minutes at room temperature. Propidium iodide staining was quantified using FACS on the Guava easyCyte (Millipore). Cell cycle analysis was performed using the FlowJo software (version 10.8.1). The statistical significance of the difference in the percentage of cells in each phase of the cell cycle was determined by multiple unpaired T-tests with Holm-Šídák multiple testing *P-value* adjustment. Each cell line was tested independently, and the technical replicates were averaged for each biological replicate.

## Acknowledgements

The authors would like to thank members of the Dunbier and Mace groups especially Abigail Burgess and Sam Jamieson, the Cancer Research UK Cambridge Institute proteomics core and bioinformatics core, especially Clive D’Santos and Kamal Kishore, and the Otago Genomics Facility. This work was funded by a project grant from the Health Research Council of New Zealand, H.D.M. was supported by University of PhD Scholarships, the University of Otago Elman Poole Traveling Scholarship and the Marjorie McCallum Travel Scholarship. The Fusion Lumos Orbitrap mass spectrometer was purchased with the support from a Wellcome Trust Multi-user Equipment Grant (Grant #108467/Z/15/Z)

## Data Availability

RNAseq data is available from the NCBI Gene Expression Omnibus (GSE233138). Raw mass spectrometry from the TRIB1 qPLEX-RIME data is available from the PRIDE database (PXD038488). All code will be made available through GitHub. All other relevant data are provided in the supplementary information or will be made available upon reasonable request.

