## Supplementary figures for "TRIB1 modulates transcriptional programming in breast cancer cells to regulate cell proliferation"

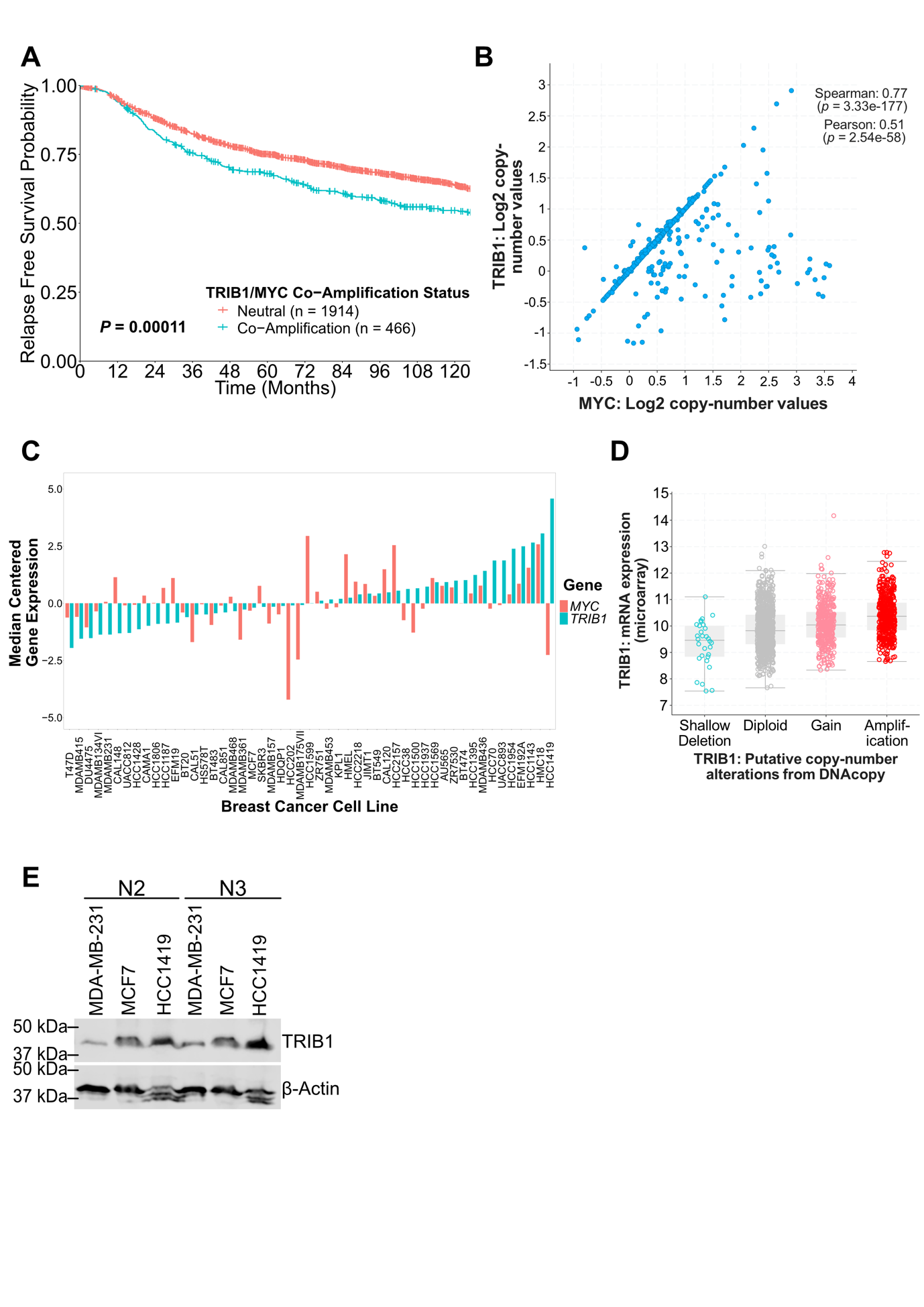

**Supplementary Figure 1: TRIB1 is co-amplified with MYC in breast cancer patient samples. A**, Kaplan-Maier survival plot of relapse free survival from the METABRIC dataset, stratified based on TRIB1/MYC co-amplification status (1–3). **B**, Correlation of estimated TRIB1 and MYC copy number from the TCGA dataset (4, 5), analysis performed using cBioPortal (2). **C**, Waterfall plot of median centered TRIB1 and MYC expression from the CCLE dataset (GSE36133) (6). **D**, TRIB1 expression in tumors from the METABRIC dataset grouped based on TRIB1 amplification status, analysis performed using cBioPortal (1–3). **E**, Western blot of TRIB1 levels in MDA-MB-231, MCF7 and HCC1419 cells after 72 hours of growth, the remaining two biological replicates used for densitometry (Fig. 1D). Each lane was loaded with 50 µg of total protein.

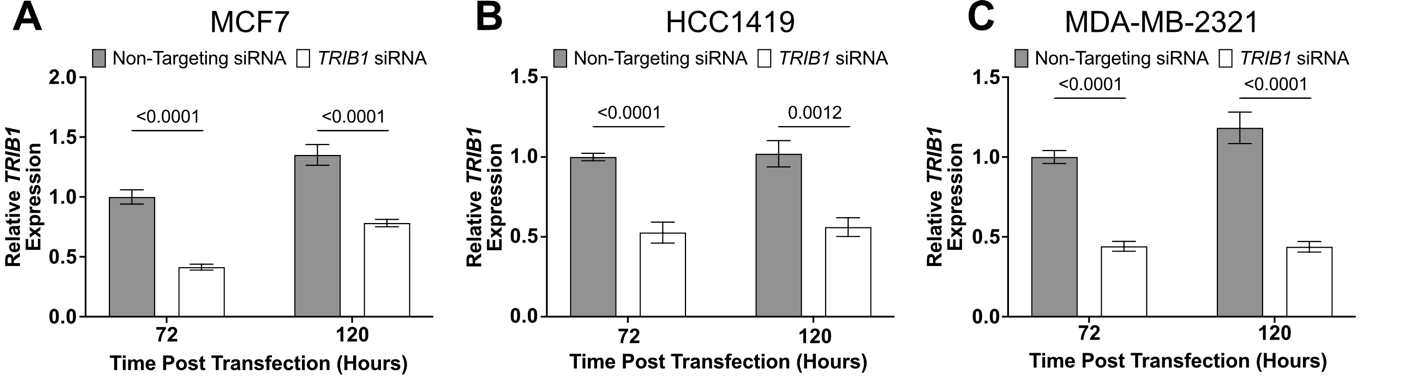

**Supplementary Figure 2: TRIB1 expression is significantly suppressed by transfection with ON-TARGET siRNA SMARTpool.** Assessment of TRIB1 expression by RT-qPCR in **A**, MCF7, **B**, HCC1419 and **C**, MDA-MB-231 cells 72 and 120 hours after transfection with TRIB1 or Non-Targeting ON-TARGET siRNA SMARTpool. All data are the mean ± SEM of three biological replicates (HCC1419 and MDA-MB-231 have two biological replicates). Significance determined independently for each cell line by multiple unpaired T-tests with Holm-Šídák multiple testing P-value correction.

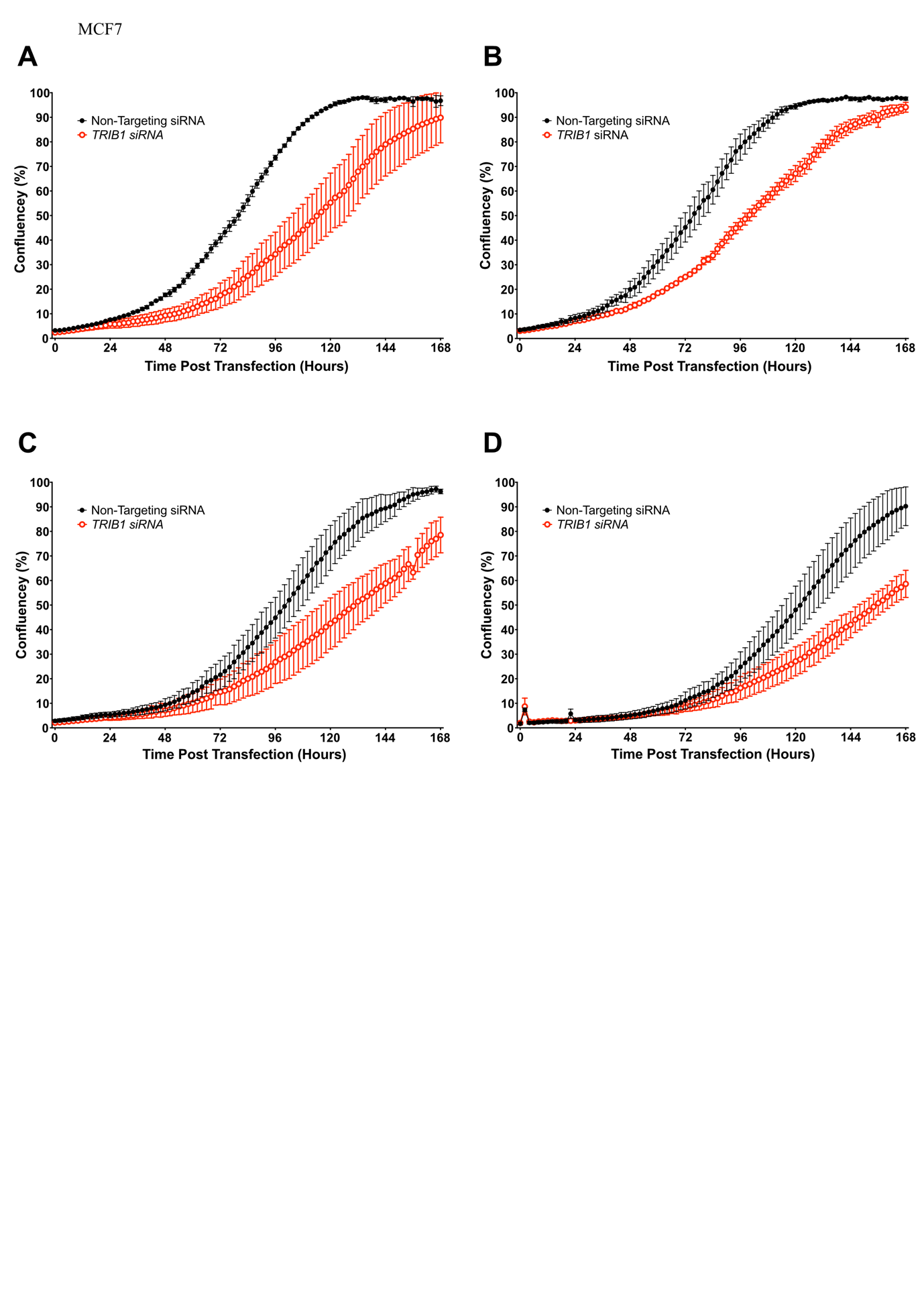

**Supplementary Figure 3: MCF7 cell proliferation is inhibited by TRIB1 knockdown with siRNA SMARTpool transfection.** Replicate **A**, One, **B**, Two, **C**, Three and **D**, Four of MCF7 cell proliferation tracking by confluency monitoring using the IncuCyte live cell imager for 168 hours post siRNA transfection. Cells were seeded at 2000 cells and transfected 24 hours later. Cell confluency was monitored with two hourly imaging, for 168 hours, with media changes 24 hours after transfection and then every 48 hours. Data are the mean confluency ± SD of technical triplicates.

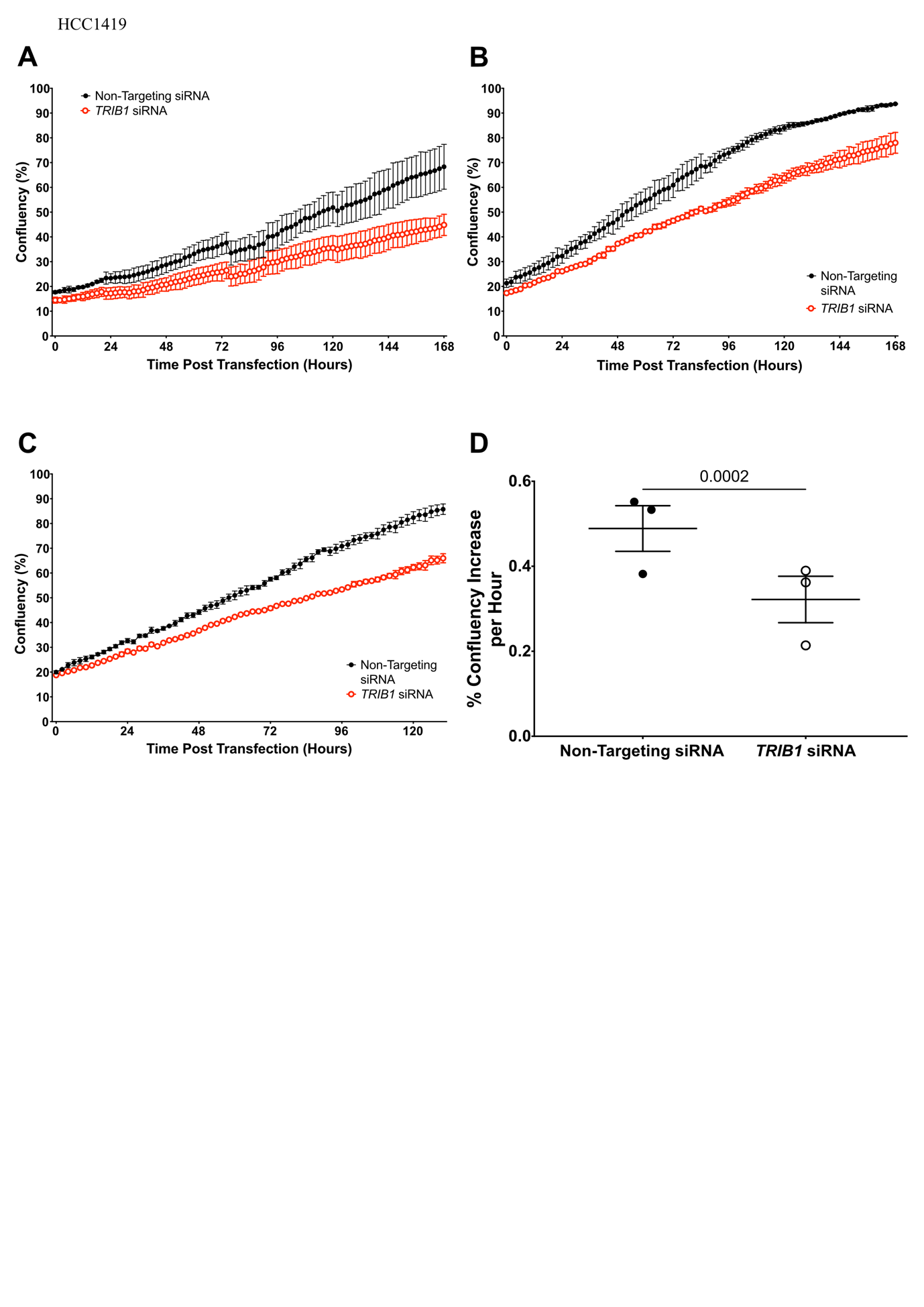

**Supplementary Figure 4: HCC1419 cell proliferation is inhibited by TRIB1 knockdown with siRNA SMARTpool transfection.** Replicate **A**, One, **B**, Two and **C**, Three of HCC1419 cell proliferation tracking by confluency monitoring using the IncuCyte live cell imager for 168 hours post siRNA transfection. Cells were seeded at 15000 cells and transfected 24 hours later. Cell confluency was monitored with two hourly imaging, for 168 hours, with media changes 24 hours after transfection and then every 48 hours. Data are the mean confluency ± SD of technical triplicates. **D**, Rate of change in confluency of HCC1419 cells after transfection with the TRIB1 or Non-Targeting ON-TARGETplus siRNA SMARTpools. Growth rate determined from IncuCyte live cell imaging confluency traces collected for 168 hours post-transfection (Supplementary Fig. 4A-C). Data are the mean ± SD of three biological replicates with significance determined by paired T-test, samples paired based on transfection/biological replicate.

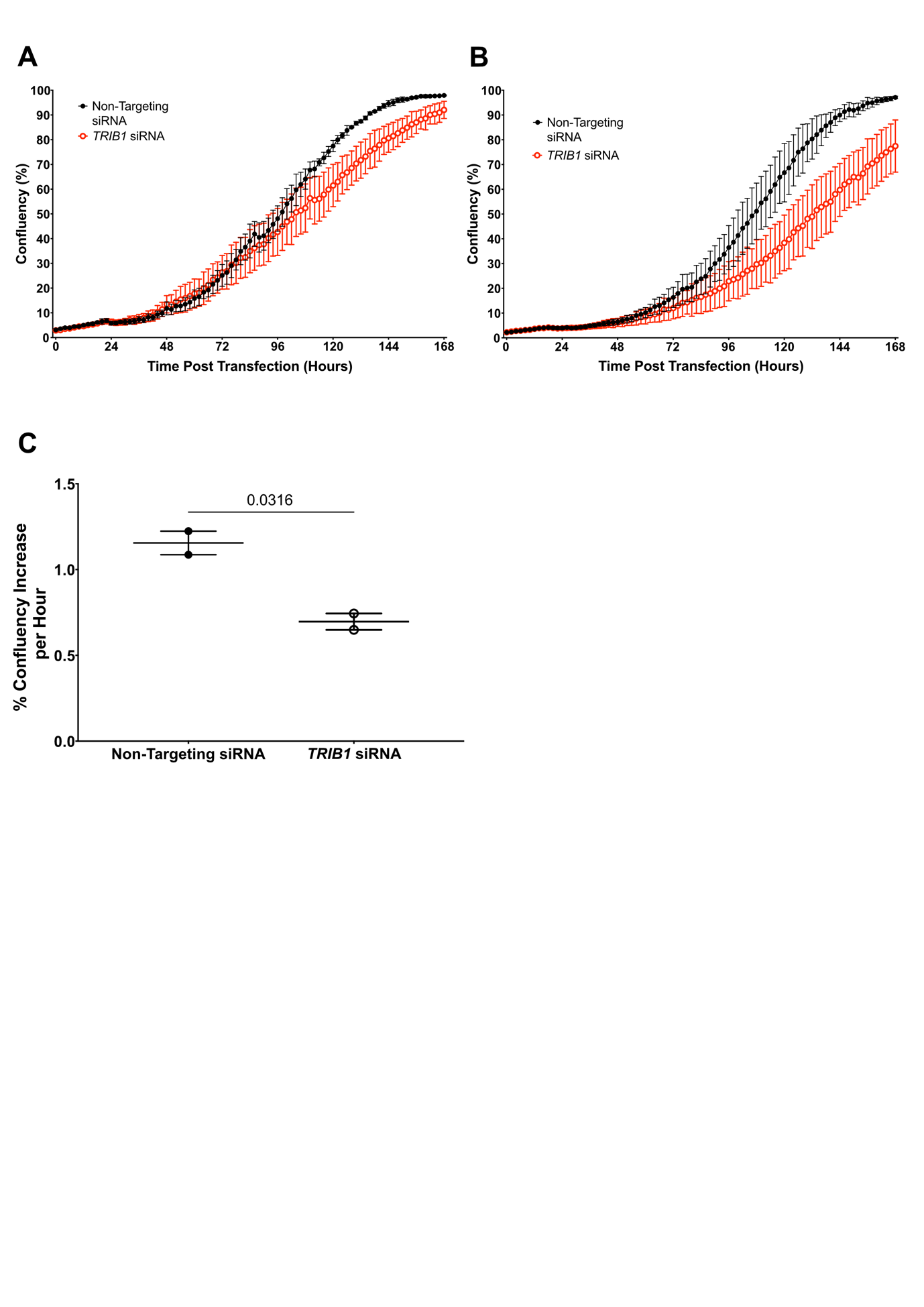

**Supplementary Figure 5: MDA-MB-231 cell proliferation is inhibited by TRIB1 knockdown with siRNA SMARTpool transfection.** Replicate **A**, One and **B**, Two of MDA-MB-231 cell proliferation tracking by confluency monitoring using the IncuCyte live cell imager for 168 hours post siRNA transfection. Cells were seeded at 2000 cells and transfected 24 hours later. Cell confluency was monitored with two hourly imaging, for 168 hours, with media changes 24 hours after transfection and then every 48 hours. Data are the mean confluency ± SD of technical triplicates. **C**, Rate of change in confluency of MDA-MB-231 cells after transfection with the TRIB1 or Non-Targeting ON-TARGETplus siRNA SMARTpools. Growth rate determined from IncuCyte live cell imaging confluency traces collected for 168 hours post-transfection (Supplementary Fig. 5A-B). Data are the mean ± SD of two biological replicates, significance determined by unpaired T-test.

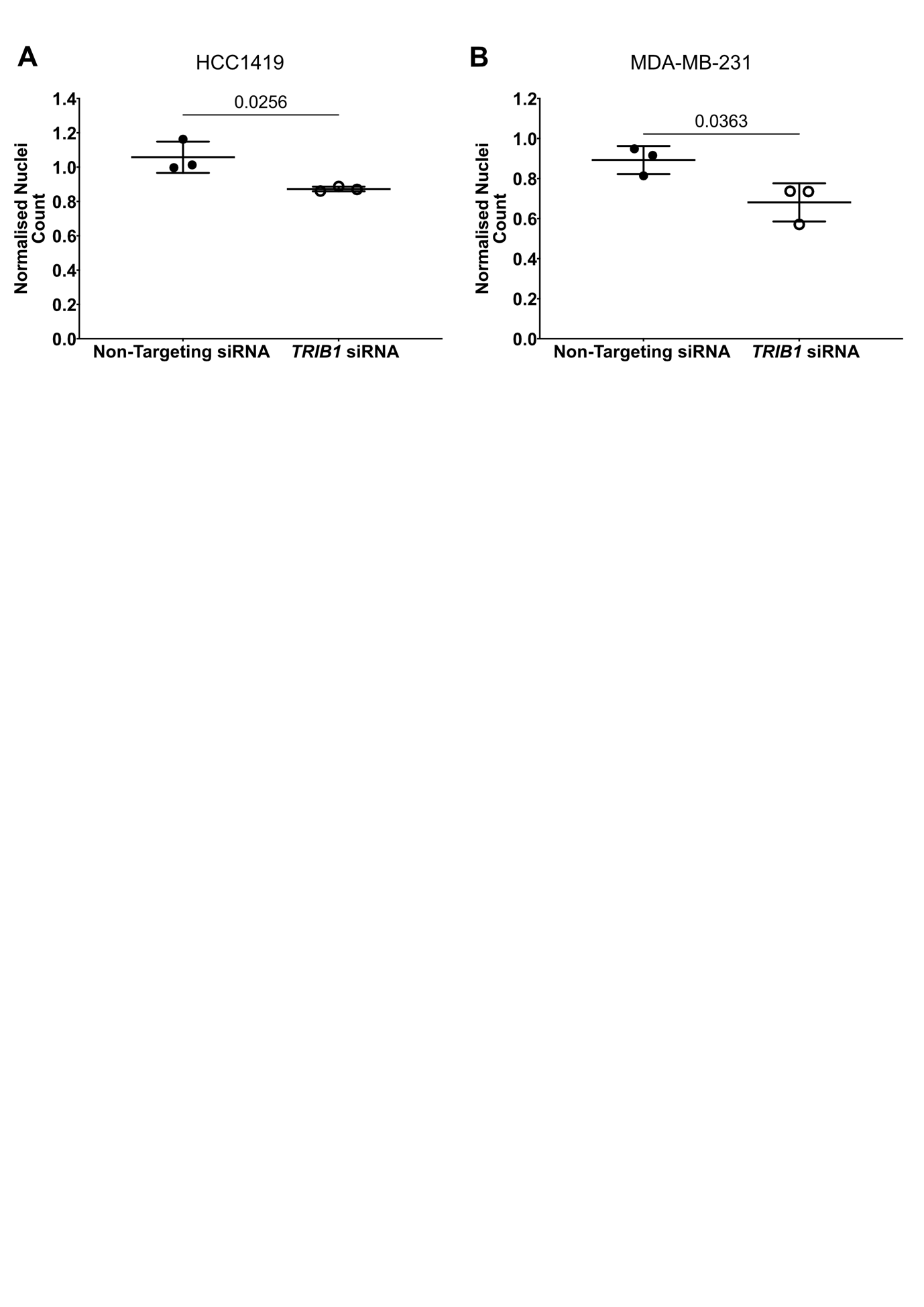

**Supplementary Figure 6: TRIB1 knockdown in HCC1419 and MDA-MB-231 inhibits proliferation resulting in decreased cell number after transfection. A**, HCC1419 nuclei count 120 hours, and **B**, MDA-MB-231 nuclei counts 72 hours after transfection with TRIB1 or Non-Targeting ON-TARGETplus siRNA SMARTpools, normalized to untreated nuclei count. Data are the mean ± SEM of three biological replicates, each containing three technical replicates. Significance determined by paired T-test, samples paired based on transfection/biological replicate.

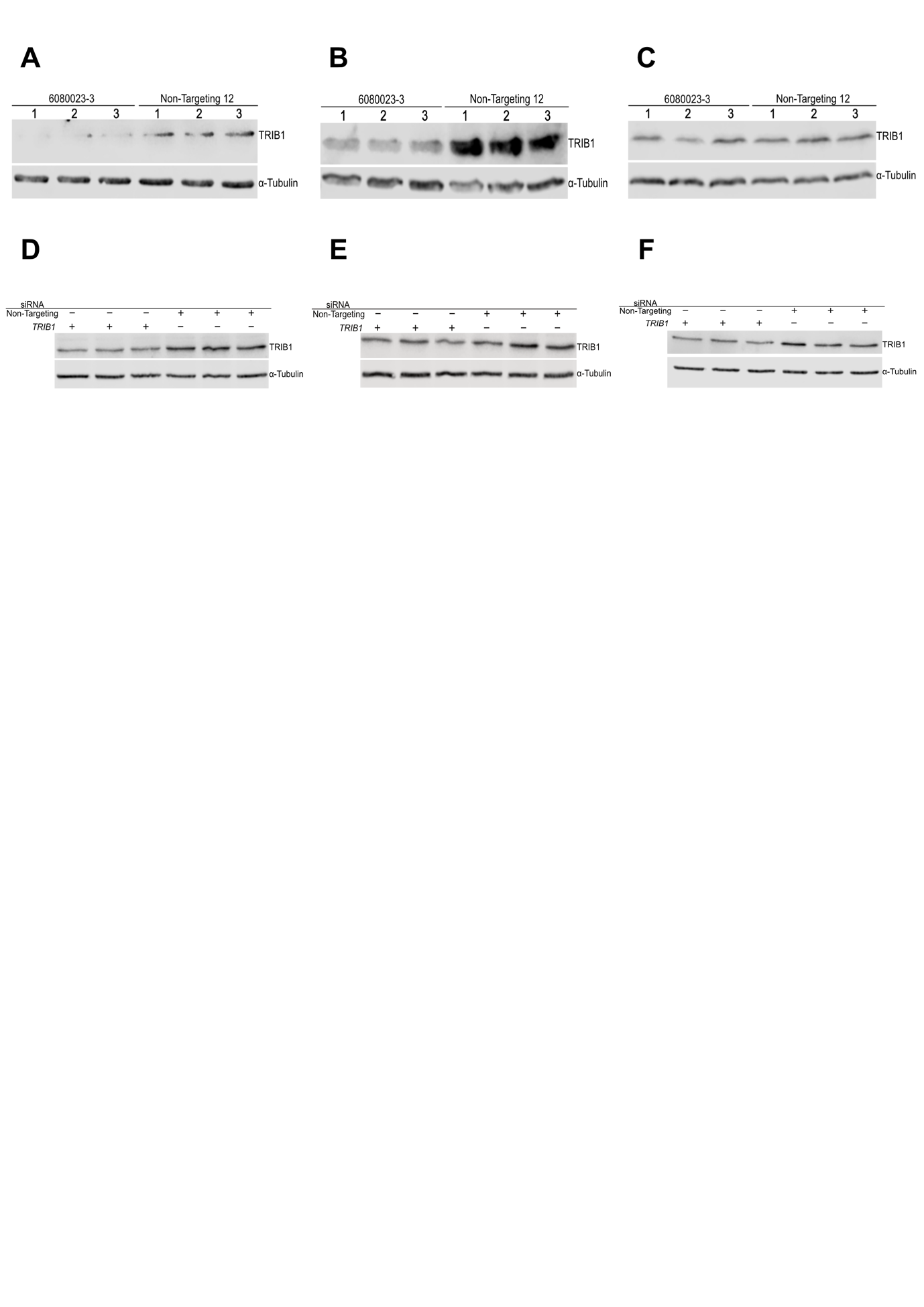

**Supplementary Figure 7: Western blots matched with RNAseq samples to confirm protein level TRIB1 knockdown. A-C,** Western blot for TRIB1 with protein lysate collected from 6080023-3 and Non-Targeting-12 MCF7 monoclones treated with 2 µg/mL doxycycline in parallel to those treated and used for RNAseq. Gel was loaded with 50 mg total protein for each time point. **D-F,** Western blot for TRIB1 with protein lysate collected from MCF7 cells treated with siRNA SMARTpools in parallel to those treated and used for RNAseq. Gel was loaded with 50 mg total protein in each lane.

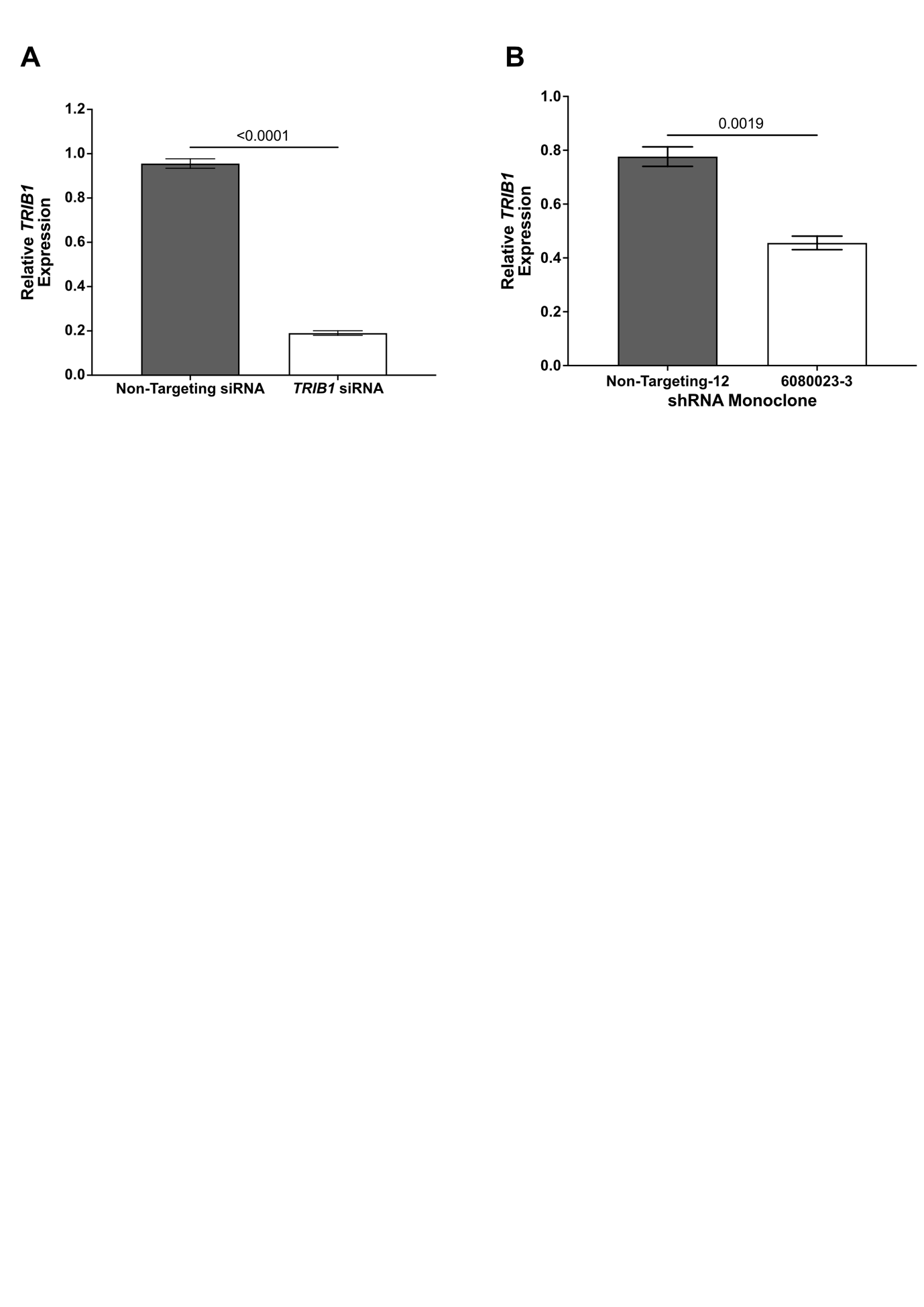

**Supplementary Figure 8: RT-qPCR evaluation of TRIB1 knockdown in samples used for RNAseq.** TRIB1 expression, as determined by RT-qPCR, in **A,** siRNA transfected MCF7 or **B**, MCF7 monoclone samples used for RNAseq analysis. Expression was normalized to the geometric mean of PUM1 and FKBP15 expression. All data is mean ± SEM of three biological replicates, each containing three technical replicates. Technical replicates were averaged for significance calculations by unpaired T-test, performed independently on the siRNA and shRNA knockdown.

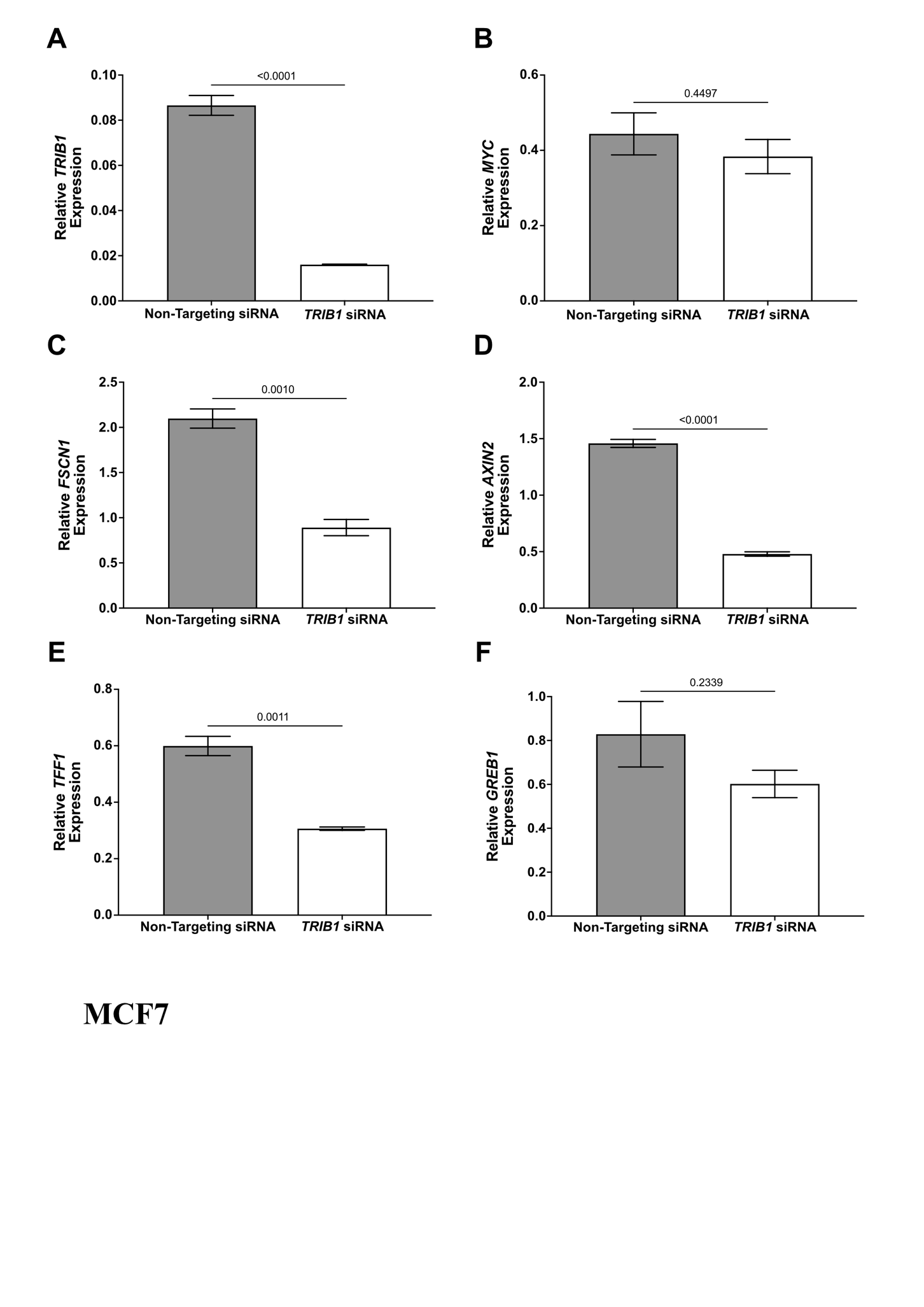

**Supplementary Figure 9: RT-qPCR evaluation gene expression in MCF7 cells after TRIB1 knockdown by siRNA SMARTpool transfection, biologically independent samples to those used for RNAseq. A,** TRIB1, **B**, MYC, **C**, FSCN1, **D**, AXIN2, **E**, TFF1 and **F**, GREB1 expression, in MCF7 cells, was determined by RT-qPCR after 48 hours of TRIB1 knockdown by siRNA SMARTpool transfection. Expression was normalized to the geometric mean of PUM1 and FKBP15 expression. All data are the mean ± SEM of three biological replicates, with the significance determined by unpaired T-test, performed independently for each gene.

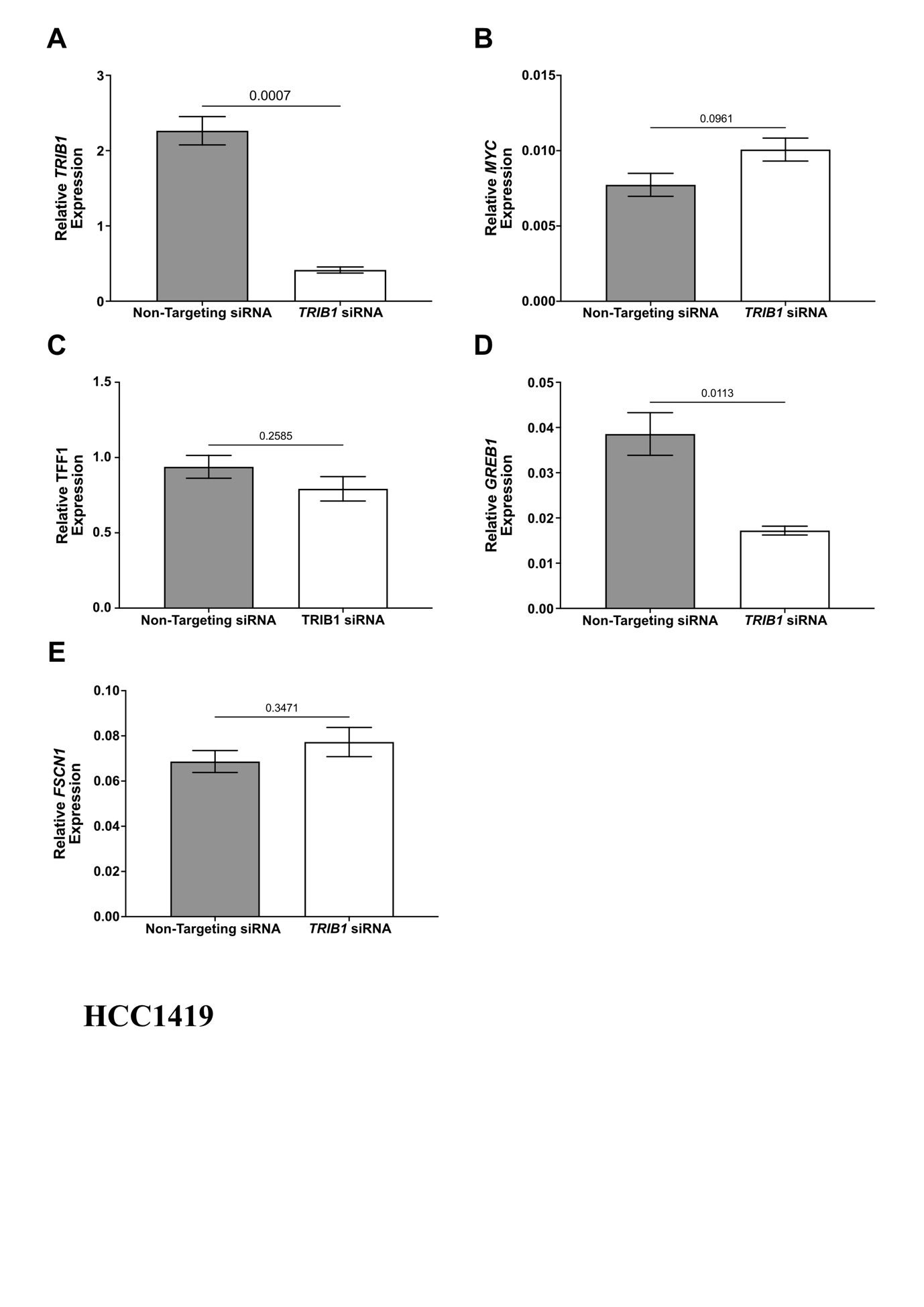

**Supplementary Figure 10: RT-qPCR evaluation of gene expression in HCC1419 cells after TRIB1 knockdown by siRNA SMARTpool transfection. A,** TRIB1, **B**, MYC, **C**, TFF1, **D**, GREB1, **E**, FSCN1 expression, in HCC1419 cells, was determined by RT-qPCR after 48 hours of TRIB1 knockdown by siRNA SMARTpool transfection. Expression was normalized to the geometric mean of PUM1 and FKBP15 expression. All data are the mean ± SEM of three biological replicates, with the significance determined by unpaired T-test, performed independently for each gene.

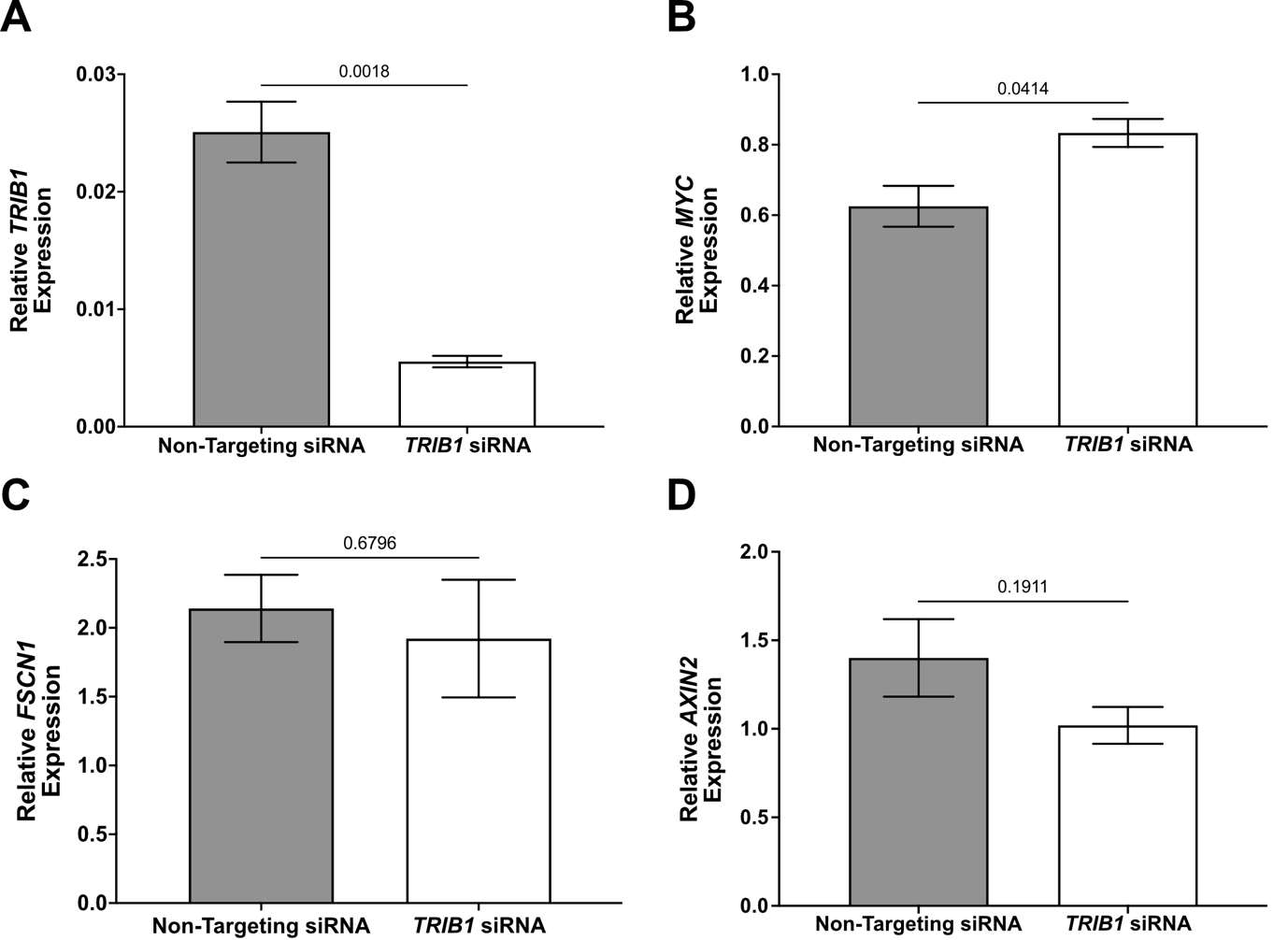

**Supplementary Figure 11: RT-qPCR evaluation gene expression in MDA-MB-231 cells after TRIB1 knockdown by siRNA SMARTpool transfection. A,** TRIB1, **B**, MYC, **C**, FSCN1, **D**, AXIN2, expression, in MDA-MB-231 cells, was determined by RT-qPCR after 48 hours of TRIB1 knockdown by siRNA SMARTpool transfection. Expression was normalized to the geometric mean of PUM1 and FKBP15 expression. All data are the mean ± SEM of three biological replicates, with the significance determined by unpaired T-test, performed independently for each gene.

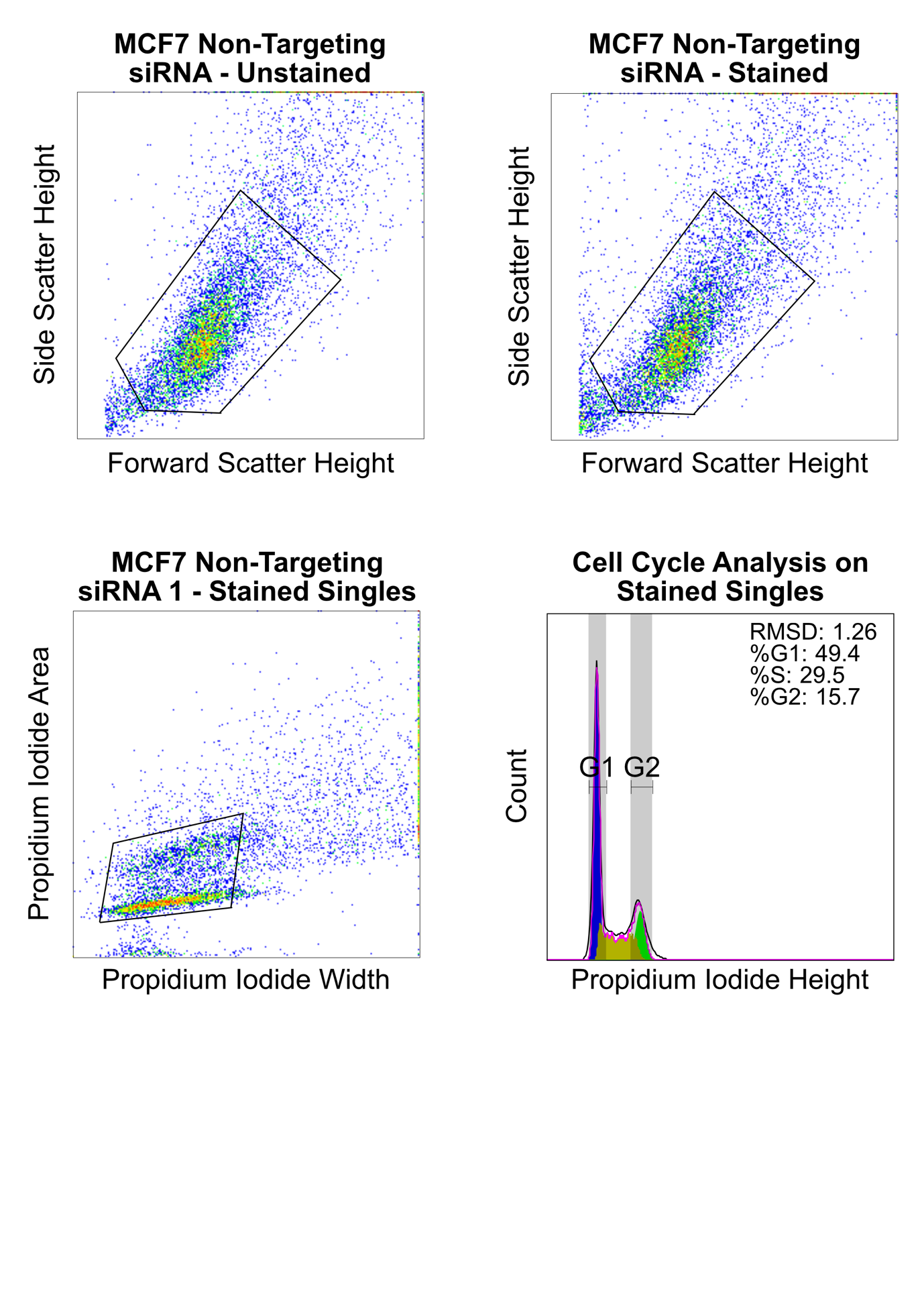

**Supplementary Figure 12: Exemplar of gating strategy for cell cycle analysis after siRNA transfection.** Cells were initially gated off the unstained non-targeting transfected control and this gate was applied to all samples. Single stained cells were gated by comparing the width and area of the Propidium Iodide signal. Cells with very wide or high area were excluded from analysis as doublets. The cell cycle distribution of the selected stained single cells was determined using the cell cycle analysis tool on FlowJo software (Version 10.8.1)

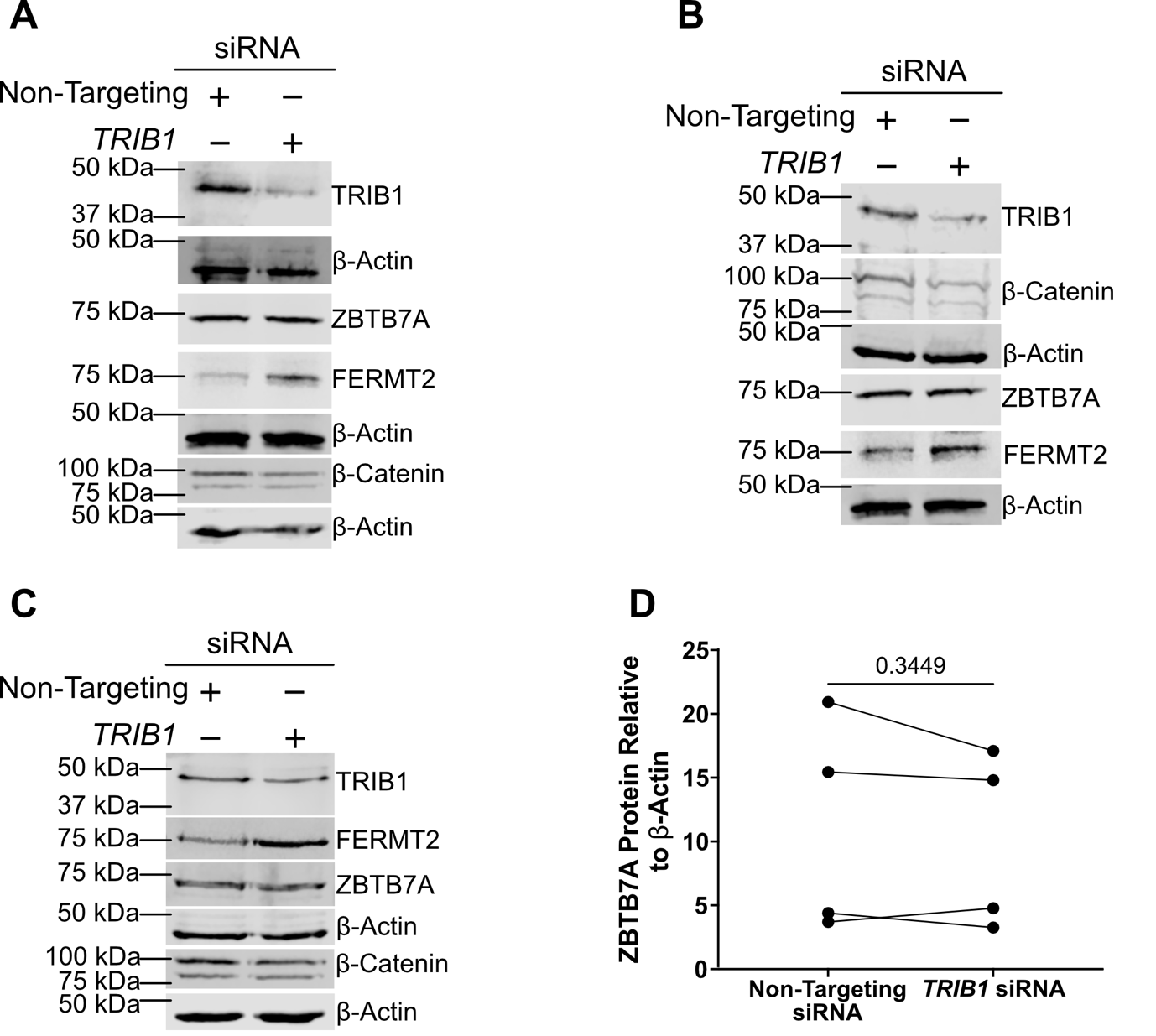

**Supplementary Figure 13: Western blot evaluation of protein levels in MCF7 cells 72 hours after transfection with TRIB1 or Non-Targeting siRNA SMARTpools. A-C,** western blots of lysate from MCF7 cells 72 hours after transfection with TRIB1 or Non-Targeting siRNA SMARTpools. Each western blot is a biological replicate. Gel loaded with 50 µg of total protein. Densitometry of ZBTB7A (determined using ImageStudioLite) normalised to β-actin, each point is a biological replicate. Significance determined by paired T-test.
